# Centriolar satellites are sites of translation of centrosomal and ciliary proteins

**DOI:** 10.1101/2024.02.22.581531

**Authors:** Claudia Pachinger, Jeroen Dobbelaere, Cornelia Rumpf-Kienzl, Shiviya Raina, Júlia Garcia-Baucells, Marina Sarantseva, Andrea Brauneis, Alexander Dammermann

## Abstract

Centriolar satellites are cytoplasmic particles found in the vicinity of centrosomes and cilia whose functional contribution to the formation of these cellular structures has long been unclear. By characterizing the main scaffolding component of satellites, PCM1 or Combover in *Drosophila*, we show that satellites are not involved in cellular trafficking as previously thought but rather act as sites for the coordinate translation of centrosomal and ciliary proteins through the interaction with a set of RNA binding proteins and proteins involved in quality control. Strikingly, the concentration of satellites near centrosomes and cilia in vertebrates is not a conserved feature and therefore dispensable for satellite function. Such coordinate synthesis may be a general feature in eukaryotic cells to facilitate protein complex formation and cellular compartmentalization.

**One-Sentence Summary:** Centriolar satellites facilitate the coordinate synthesis of centrosomal and ciliary proteins.

## Results and Discussion

Centriolar satellites are non-membranous particles concentrated in the vicinity of centrosomes and the ciliary base in vertebrate cells (*1, 2*). Work over the past two decades has linked numerous proteins to satellites, including proteins otherwise localized to centrioles, the pericentriolar material (PCM) or cilia (*3–5*). However, just a single protein, pericentriolar material 1 (PCM1), has been found to be essential for their formation, suggesting that it acts as their main scaffolding component (*6*). PCM1 depletion not only eliminates satellites but also impairs the accumulation of centrosomal and ciliary proteins, thereby affecting processes from centriole assembly to centrosomal microtubule anchorage and ciliary trafficking (*6–8*). This, together with the reported dynein motor-dependent movement of satellites to centrosomes (*9*) has led to the idea of their functioning as transport modules delivering proteins for centrosome and cilium biogenesis (*1, 2, 6*) (Fig. 1A).

**Fig. 1.**
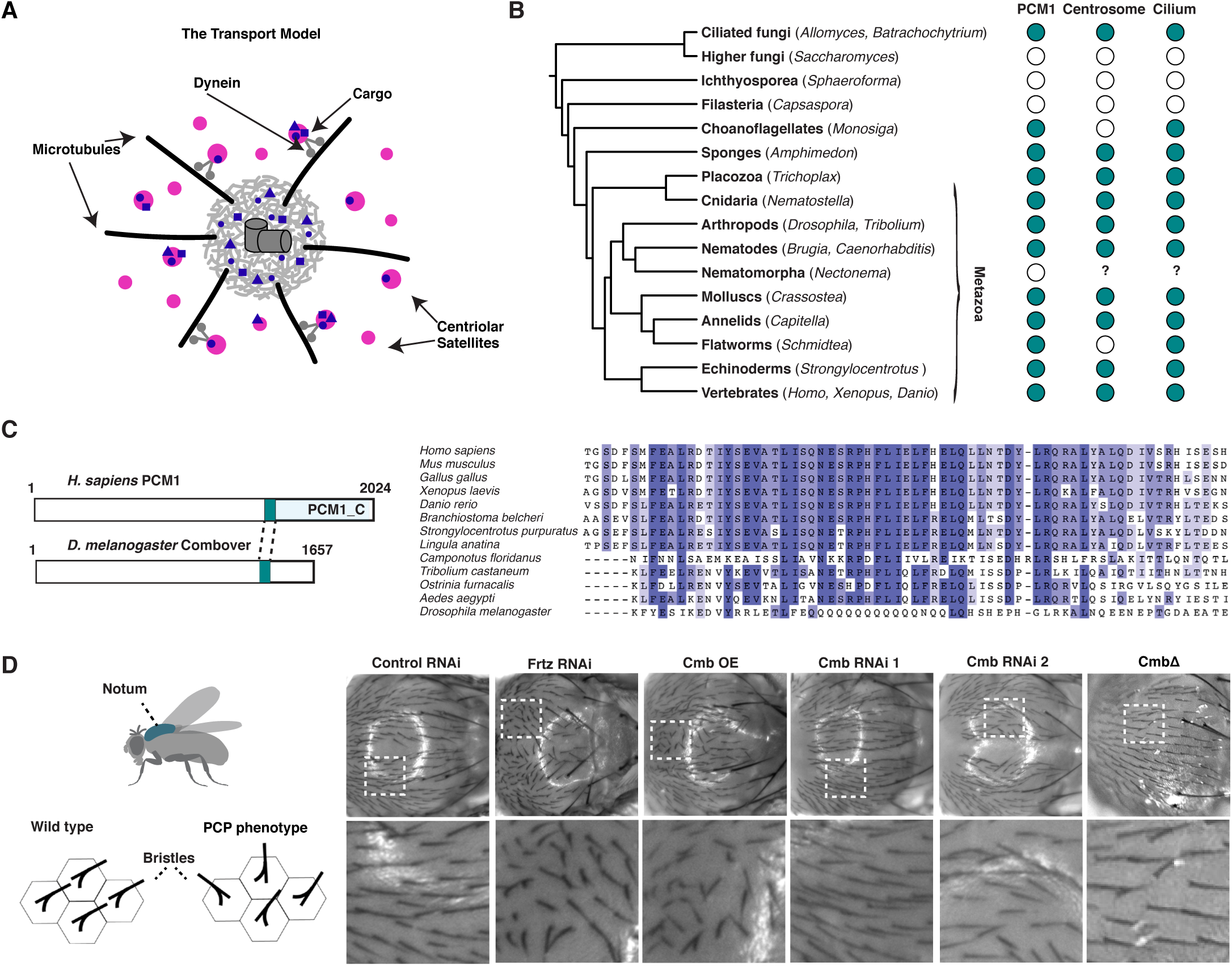
The satellite scaffolding component PCM1 is conserved beyond vertebrates. (**A**) Schematic representation of the transport model of centriolar satellite (magenta) function as mediators of dynein-dependent recruitment of centrosomal and ciliary client proteins (blue) for centrosome/cilium biogenesis. (**B**) Reciprocal BLAST analysis reveals the presence of PCM1 orthologs across opisthokonts, correlating with the reported presence of centriole-organized centrosomes and cilia (*58, 59*). See also fig. S1A. (**C**) Multiple sequence alignment of conserved C-terminus (part of pfam15717) of selected PCM1 orthologs. Note that *Drosophila* Combover (Cmb) is highly divergent. (**D**) Overexpression but not RNAi-mediated depletion or mutation of Cmb results in a planar cell polarity (PCP) phenotype, seen by misalignment of bristles in the fly notum. Tissue-specific RNAi performed using the Pannier-GAL4 driver, with the PCP effector Fritz (*24*) as positive control.

Yet, a number of observations have called into question the transport model. While satellite perturbations are highly pleiotropic and affect nearly every aspect of centrosome and cilium biology, phenotypes are not as severe as might be expected considering their wide range of centrosomal and ciliary clients. Thus, while mouse mutants in *PLK4/SAK* (*10*), *C2CD3* (*11*) and *PCNT* (*12*), all encoding satellite proteins, exhibit fully penetrant prenatal lethality, *PCM1* mutants are born at Mendelian frequencies, albeit with significant postnatal lethality and exhibiting a variety of defects indicative of ciliary dysfunction (*13*). Centrosome composition is also largely unaffected (*4, 5*). The whole, then, is not greater than the sum of its parts. Also, for trafficking modules evidence for directed motion of satellites is surprisingly sparse, with live cell imaging in cultured cells revealing PCM1 particles moving largely by diffusion (*14*). At the same time satellite distribution is highly variable, responding to a variety of environmental factors and cellular stresses (*15, 16*) as well as cell cycle state, with a marked disassembly in mitotic cells (*6*). These changes may, however, be driven not by motor-mediated movement, but by liquid-liquid phase separation/dissolution of PCM1 condensates (*17, 18*).

Here, we re-examine satellite function, building on the identification of a *Drosophila* ortholog of PCM1. Defining features conserved across metazoan evolution, we identify a role for satellites not in protein trafficking, but in the coordinate synthesis of centrosomal and ciliary proteins, a paradigm shift we suggest reconciles these disparate observations.

## The satellite scaffolding component PCM1 is conserved beyond vertebrates

While invertebrate model organisms including *C. elegans* and *Drosophila melanogaster* have made significant contributions to our understanding of centrosome and cilium biogenesis, work on satellites has so far focused exclusively on their role in vertebrates and in particular vertebrate cultured cells. Indeed, centriolar satellites have long been thought to be unique to vertebrates. Yet, a computational study identified putative orthologs of PCM1 in both *C. elegans* and *Drosophila* (*19*), suggesting the phylogenetic distribution of PCM1 and satellites may be wider than currently appreciated. Reciprocal BLAST analysis confirmed this identification and revealed further orthologs across opisthokonts, including in choanoflagellates and ciliated fungi (Blastocladiales, chytrids), though not in higher fungi, in which centrioles and cilia have been secondarily lost (Fig. 1B, see also fig. S1A). PCM1 orthologs could also not be identified in representatives of the parasitic phylum Nematomorpha (*20*), which lack most ciliary genes. In contrast, PCM1 is conserved in the planarian *Schmidtea mediterranea*, which lacks centrosomes but retains centrioles and cilia (*21*). This pattern of inheritance and loss is consistent with a functional association with centriole-based structures conserved across >1000 million years (*22*) of opisthokont evolution.

We identified the *Drosophila* ortholog of PCM1, previously named Combover or Cmb (*23*), in the course of establishing TurboID in flies as a proximity interactor of the centriolar structural component Sas4 in S2 cells (fig. S1B). Primary sequence homology to the human protein, largely restricted in insects to the more conserved C-terminus, is extremely low (Fig. 1C). We were therefore interested to determine to what extent the functions ascribed to vertebrate PCM1 are conserved in *Drosophila*. Unlike the putative *C. elegans* ortholog, Cmb is not entirely uncharacterized, having been linked to planar cell polarity (PCP), with overexpression of the protein resulting in the formation of multiple hair cells in the wing (*23*), a phenotype associated with perturbation of PCP effector genes such as Fritz (Frtz), Fuzzy (Fy) and Inturned (In) (*24*). We observed similar defects in the orientation of bristles upon overexpression of Cmb in the notum of the fly, another signature PCP effector phenotype (Figs. 1D, S1C,D). However, neither acute depletion by tissue-specific RNAi nor complete loss of the protein in *Cmb* mutants resulted in any observable PCP phenotype. The functional significance of this link therefore remains unclear.

## *Drosophila* PCM1/Cmb is required for proper centrosome and cilium function

While *Cmb* mutants appear morphologically wild type, behaviorally they are clearly not. The first noticeable defect is abnormal wing posture, indicative of impaired mechanosensation (*25*) (fig. S2A) involving the chordotonal neurons, a ciliated tissue in the fly (Fig. 2A). Consistent with this, negative geotaxis in adult flies (*26*) was found to be strongly impaired (Fig. 2B). Mutants were also fully male (though not female) infertile (Figs. 2C, S2B). Both types of defect could be reproduced by tissue-specific RNAi and rescued by introduction of a GFP transgene, confirming specificity of the mutant phenotype and functionality of the GFP transgene. Neither phenotype is shared by PCP genes, but both are highly reminiscent of what is observed following perturbation of centriolar or ciliary components (*27, 28*). A closer examination of ciliary morphology in the chordotonal neurons responsible for mechanosensation in the animals’ legs by DIC and immunofluorescence microscopy as well as ultrastructural analysis revealed only minor defects (fig. S2C-E). In contrast, spermatogenesis was more markedly affected, with a dissection of the testes of adult males revealing weakly motile sperm incapable of reaching the seminal vesicle (fig. 2F). Transmission electron microscopy showed a significant proportion of broken axonemes and missing axonemal microtubule doublets (Fig. 2D). Cysts furthermore contained fewer than 64 flagella, indicative of a failure of axoneme extension or prior cell division defects (Fig. 2E). Examining the process of spermatogenesis by staining isolated testes to visualize nuclear morphology and the actin cytoskeleton revealed no apparent defects at early stages of differentiation until the formation of actin cones during individualization, a process that was highly defective in *Cmb* mutants (fig. S2G).

**Fig. 2.**
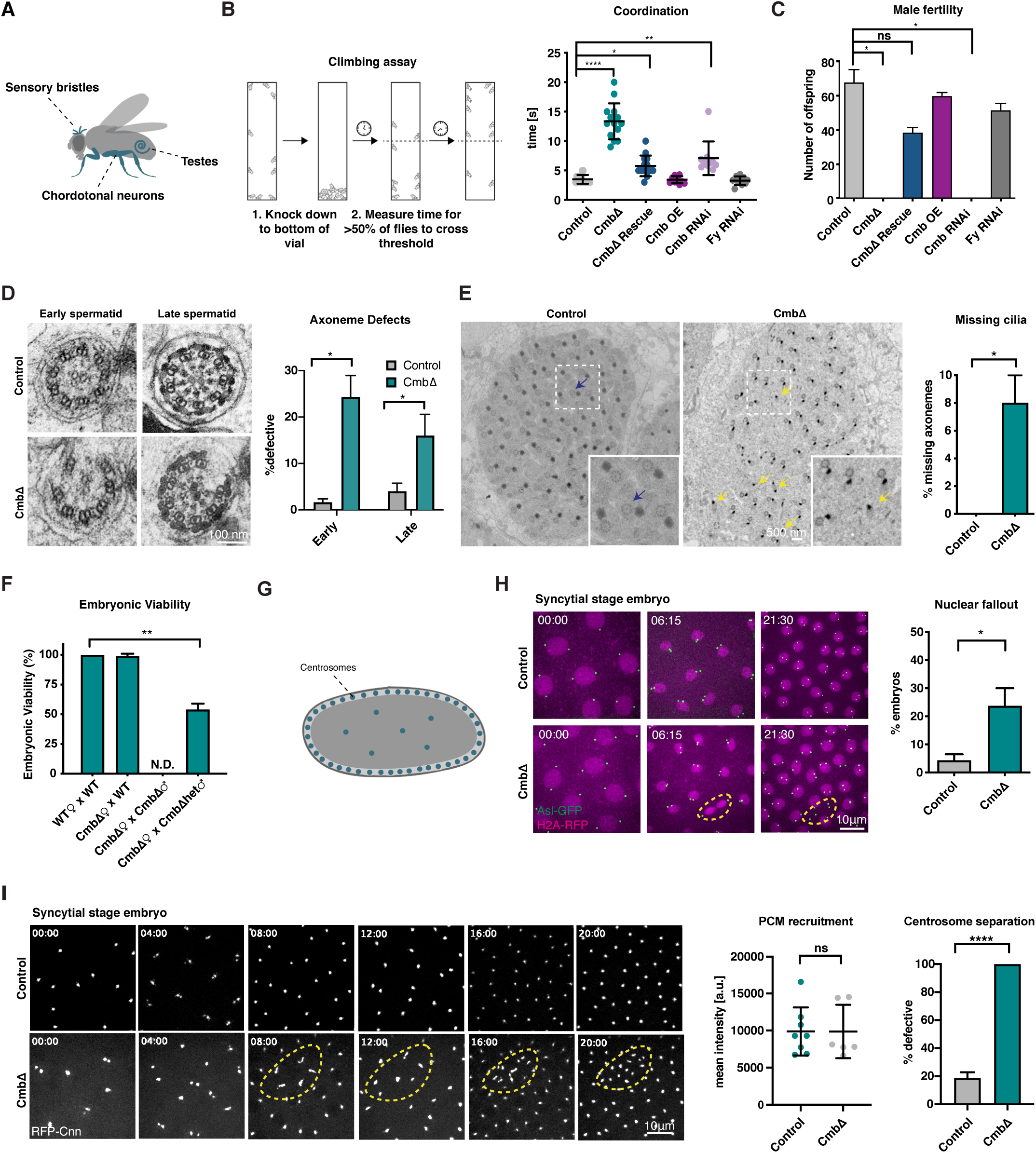
Combover is required for ciliogenesis and proper cell division. (**A**) Schematic of ciliated tissues used for phenotypic analysis. Primary cilia are found in sensory bristles and chordotonal neurons, while motile cilia/flagella are found in testes (*60*). (**B**) Climbing assay used to assess defects in mechanosensation. Cmb mutant flies are severely uncoordinated, a phenotype rescued by expression of Cmb-GFP. PCP flies (Fy RNAi) show no detectable phenotype. N>10 flies per condition. Kruskal-Wallis test with Dunn’s multiple comparisons test was performed; *P<0.05, **P< 0.01, ***P< 0.001 (**C**) Male fertility scored by crossing individual males with WT virgins and assessing the number of offspring. Cmb RNAi/mutant flies are fully male infertile. N=3 single males each crossed to 4 virgin females per condition. Kruskal-Wallis test with Dunn’s multiple comparisons test was performed; *P< 0.05. (**D**, **E**) TEM analysis of sperm axonemes in control and Cmb mutant testes. Cross-sectional views reveal missing axonemal doublets and fragmented axonemes (D) as well as overall lower number of cilia (number per cyst < 64). N=38 cysts control, 38 cysts Cmb mutant. Student’s t-test used to assess statistical significance; *P<0.05. (**F**) Embryonic viability test shows lethality in 50% of offspring of Cmb mutant females with heterozygous mutant males (viability could not be assessed for homozygous males due to their failure to mate and fertilize oocytes). N=2 virgin females crossed to one male per condition. Student t-test used to asses statistical significance; **P<0.01. (**G**) Schematic of *Drosophila* syncytial-stage embryo showing synchronous nuclear divisions occurring close to the egg surface. (**H**) Live imaging of control and Cmb mutant early embryos (nuclear cycle 12) expressing the centriole marker Asl-GFP and H2A-RFP. Time shown in min:s. Cmb mutants show lagging chromosomes and increased frequency of nuclear fallout (orange dashed circle). N=6 control, 8 Cmb mutant embryos. Student’s t-test was used to assess statistical significance; *P< 0.05. (**I**) Live imaging of control and Cmb mutant early embryos (nuclear cycle 12) expressing the centrosome marker RFP-Cnn. Time shown in min:s. Cmb mutants show defects in centrosome separation but not PCM recruitment. PCM recruitment is unaffected in Cmb mutants. Cnn centrosome intensity measured at nuclear envelope breakdown (NEBD) in nuclear cycle 12. Dots represent average intensity of all centrosomes in a single embryo. N=8 control, 6 Cmb mutant embryos. Mann-Whitney test used to assess statistical significance. Centrosome separation assessed at nuclear cycle 12 and 13. N=16 control, 12 Cmb mutant embryos. Mann-Whitney test used to assess statistical significance; ****P< 0.0001.

*Cmb* mutants, then, display clear defects in cilium biogenesis and function. However, ciliary defects cannot explain the fully penetrant parental effect embryonic lethality of *Cmb* mutants. Thus, while homozygous mutant mothers are viable if uncoordinated, the offspring of these mothers and heterozygous males (homozygous males being unable to mate and fertilize oocytes) exhibit 50% lethality, with survivors invariably found to be those 50% expected to carry a wild-type allele (Fig. 2F). Examination of syncytial stage embryos resulting from such a cross expressing markers for centrioles (Asl, green) and chromosomes (Histone H2A, red) revealed a high degree of chromosome missegregation followed by nuclear fallout (Fig. 2G, H), a quality control mechanism that internalizes faulty nuclei to the embryo interior to prevent them from contributing to further development (*29*). Centrioles and centrosomes are famously dispensable for later stages of development and morphogenesis in the fly (*30*). However, the same is not true for early embryogenesis, which in flies as in vertebrates (*10, 31*) depends on centrosomal microtubule-organizing center activity to sustain spindle assembly during their rapid mitotic cell division cycles (*32*). Time-lapse imaging of Asl revealed no evidence for centriole duplication defects in *Cmb* mutants, nor was mitotic PCM assembly significantly affected based on analysis of the PCM scaffolding component Cnn (Fig. 2H, I). However, embryos displayed a frequent failure of centrosome separation that could explain the mitotic defects observed in *Cmb* mutants (Fig. 2I). In summary, then, Cmb is dispensable for planar cell polarity, but like PCM1 in vertebrates (*13*) is required for proper centrosome and cilium function and hence organismal viability and fertility.

## Satellites are conserved in *Drosophila* but do not concentrate near centrosomes or cilia

A defining characteristic of centriolar satellites in vertebrates is their concentration near centrosomes and the base of cilia, as well as other non-centrosomal MTOCs in differentiated cells (*33*) (Fig. 3A). This behavior is consistent with their originally ascribed role in delivery of proteins required for centrosome and cilium function. In stark contrast, while *Drosophila* Cmb forms cytoplasmic foci like vertebrate PCM1, these foci show no discernible enrichment near centrosomes or cilia in S2 cultured cells or any fly tissue (Fig. 3B, C), although Cmb is occasionally observed at centrosomes in a small fraction of cells (fig. S3A). This is true for both rescuing GFP transgene and endogenous Cmb, detected using a polyclonal antibody raised against the fly protein (fig. S3B). There is furthermore little sign of directed trafficking, with Cmb particles moving largely by diffusion or entirely stationary (Fig. 3D-H).

**Fig. 3.**
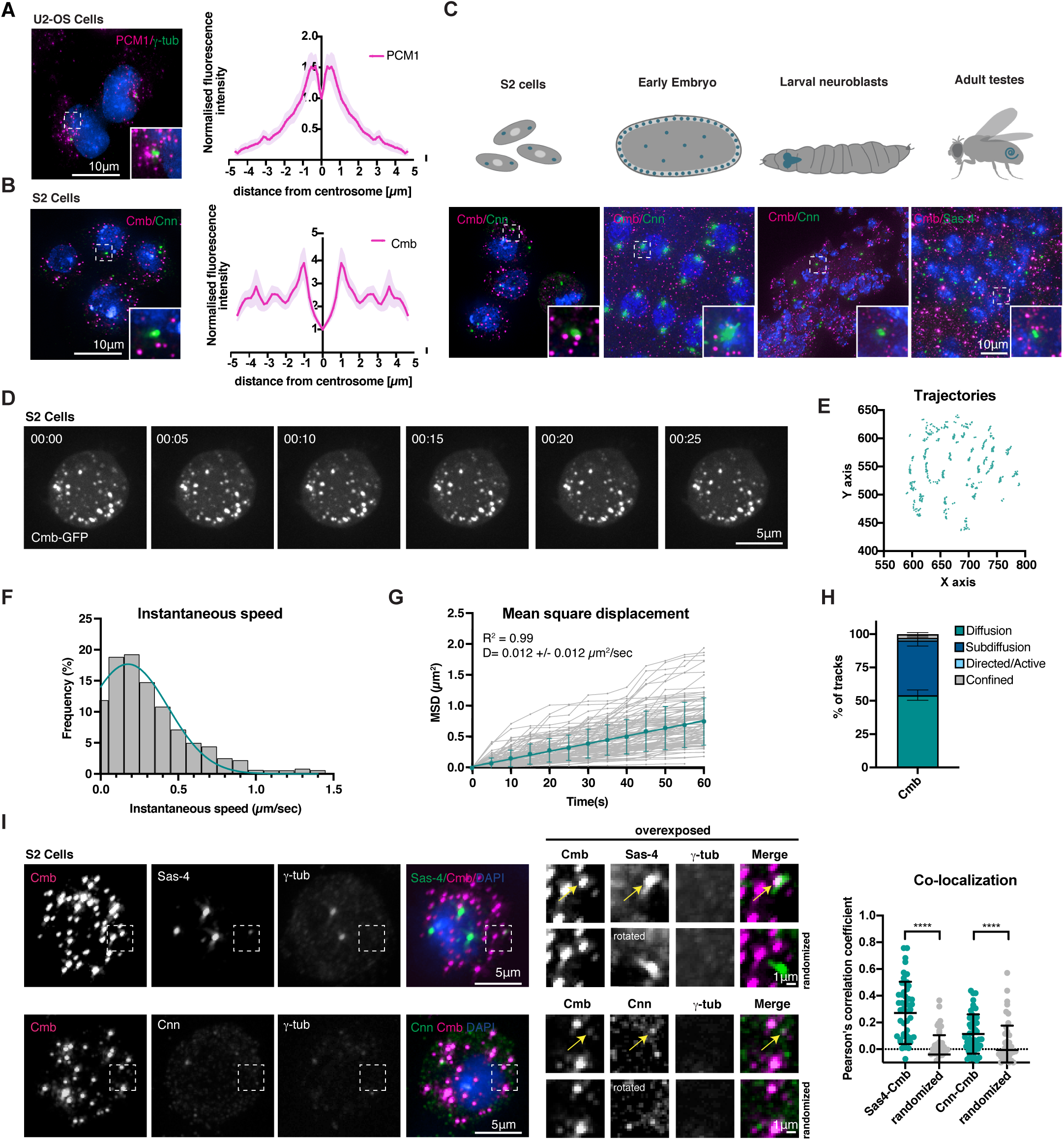
Centriolar satellites are conserved in *Drosophila* but do not concentrate near centrosomes or cilia, nor move in a directed manner. (**A**) PCM1 concentrates in the vicinity of centrosomes (marked with γ-tubulin) in vertebrate cells. Radial profile of the normalized distribution of PCM1 (N=27 centrosomes). (**B**) Cmb localization in *Drosophila* S2 cells. Unlike PCM1, Cmb does not concentrate near the centrosome (marked with Cnn), but is found throughout the cytoplasm. N=39 centrosomes. (**C**) Top: Schematics showing cells and tissues used to examine Cmb localization/dynamics. Bottom: Immunofluorescence micrographs show Cmb localizing to cytoplasmic foci in all tissues of the fly. Sas4/Cnn used to visualize centrioles/centrosomes. (**D**) Time-lapse recording of *Drosophila* S2 cell expressing Cmb-GFP. Time is shown in min:s. Cmb particles display little movement. (**E**) Corresponding trajectory analysis showing position of individual particles over time. (**F**) Histogram plotting frequency of instantaneous speed (µm/sec) of all particles tracked in 8 different cells. N=140 particles. The majority of particles moves at slow speeds (0.2-0.3µm/sec), consistent with diffusion. (**G**) Mean-Squared-Displacement (MSD) as a function of time. Trajectories with at least 11 frames from 8 cells were analyzed. All tracks analyzed shown in grey (N=99 tracks). Weighted mean square displacement (mean +/-SD) of all diffusive particles follows a linear fit (turquoise line), reflecting overall Brownian diffusion. (**H**) Stacked column plot (mean +/-SEM) showing all tracks analyzed in (G) assigned to different categories as described in Materials and Methods. The majority of tracks analyzed display Brownian diffusion (54.2% +/-3.9) or subdiffusion/anomalous diffusion (41.1% +/-4.3). Only a minor fraction displays directed/active (1.8% +/-0.7) or confined movement (2.9 % +/-1.1). (**I**) Centriolar (Sas-4) and centrosomal (Cnn) proteins co-localize with Cmb on cytoplasmic foci, although immunofluorescence signal is weak compared to that at centrosomes (marked with γ-tubulin). Colocalization of Cmb with Sas-4 and Cnn assessed by Pearson’s correlation coefficient. Randomized control has one of the channels rotated 90°. Centrosomes excluded from analysis. N=50 cells per condition. Mann-Whitney test used to test statistical significance; ****P< 0.0001. See also fig. S3C,D.

If Cmb is not enriched at centrosomes, how could it have been identified as a proximity interactor of Sas4 or contribute to centrosome and cilium function? Centriolar satellites have not previously been reported in *Drosophila*. However, it is worth bearing in mind that satellites in vertebrates were originally identified as electron-dense particles found in the vicinity of centrosomes (*34*), a property Cmb foci do not share. Furthermore, aside from PCM1 most satellite proteins primarily localize to centrosomes or cilia, with only a minor population on satellites, such that new constituents are usually identified by co-localization with PCM1 (*3, 6*). With this in mind, we re-evaluated centrosomal protein localization in *Drosophila* S2 cells and primary spermatocytes using a panel of antibodies to centriolar and PCM proteins in conjunction with Cmb. While the majority of each of these proteins indeed localizes to centrosomes, in many cases faint cytoplasmic foci could be identified and these foci co-localized with Cmb to an extent similar to what has been reported for satellite proteins in vertebrates (Figs. 3I, S3C-E) (*4*). One notable exception is the microtubule nucleator γ-tubulin, which is one of the few centrosomal proteins not found on satellites and only weakly and indirectly affected in satellite perturbations (*3–6*). Collectively, these results indicate that satellites are conserved in *Drosophila*, although they do not concentrate in the vicinity of centrosomes or cilia.

## Satellites are associated with cytoplasmic protein synthesis in *Drosophila*

In an effort to better understand satellite function in *Drosophila*, we performed proximity interaction analysis on Cmb by conventional, direct, TurboID (*35*) in S2 cells (Fig. 4A) and indirect, GFP-nanobody-targeted, TurboID (*36*) in fly testes (Fig. 4B). For the former analysis, we further fractionated cellular extracts to distinguish between interactions occurring in the general cytoplasm and (detergent-insoluble) cytoskeleton. No marked differences between the two cellular contexts were observed (figs. S4A,B). However, both preparations yielded numerous centrosomal protein proximity interactors as well as other proteins previously linked to PCM1 in vertebrates, such as the ubiquitin ligase MIB1 (*16*) and the deubiquitinase CYLD (*37*). Overall, there was a striking degree of overlap with the proteome of the centrosome-cilium interface previously defined in vertebrates, with 122 of the 412 Cmb proximity interactors conserved in humans (30%) also found amongst the high confidence interactors identified by BioID performed on 58 centriole, satellite, and ciliary transition zone proteins (*3*) (Fig. 4C) and a lesser but still significant overlap with the centriolar satellite proteome defined by (*4*) and (*5*) (fig. S4C). Largely missing were ciliary proteins, S2 cells being a non-ciliated cell type. Such proteins were, however, identified in fly testes, including components of the ciliary motility machinery and axonemal dynein assembly factors (DNAAFs) (Fig. 4B,C). IFT and BBS proteins do not contribute to cilium biogenesis in *Drosophila* sperm (*28, 38*) and were not detected.

**Fig. 4.**
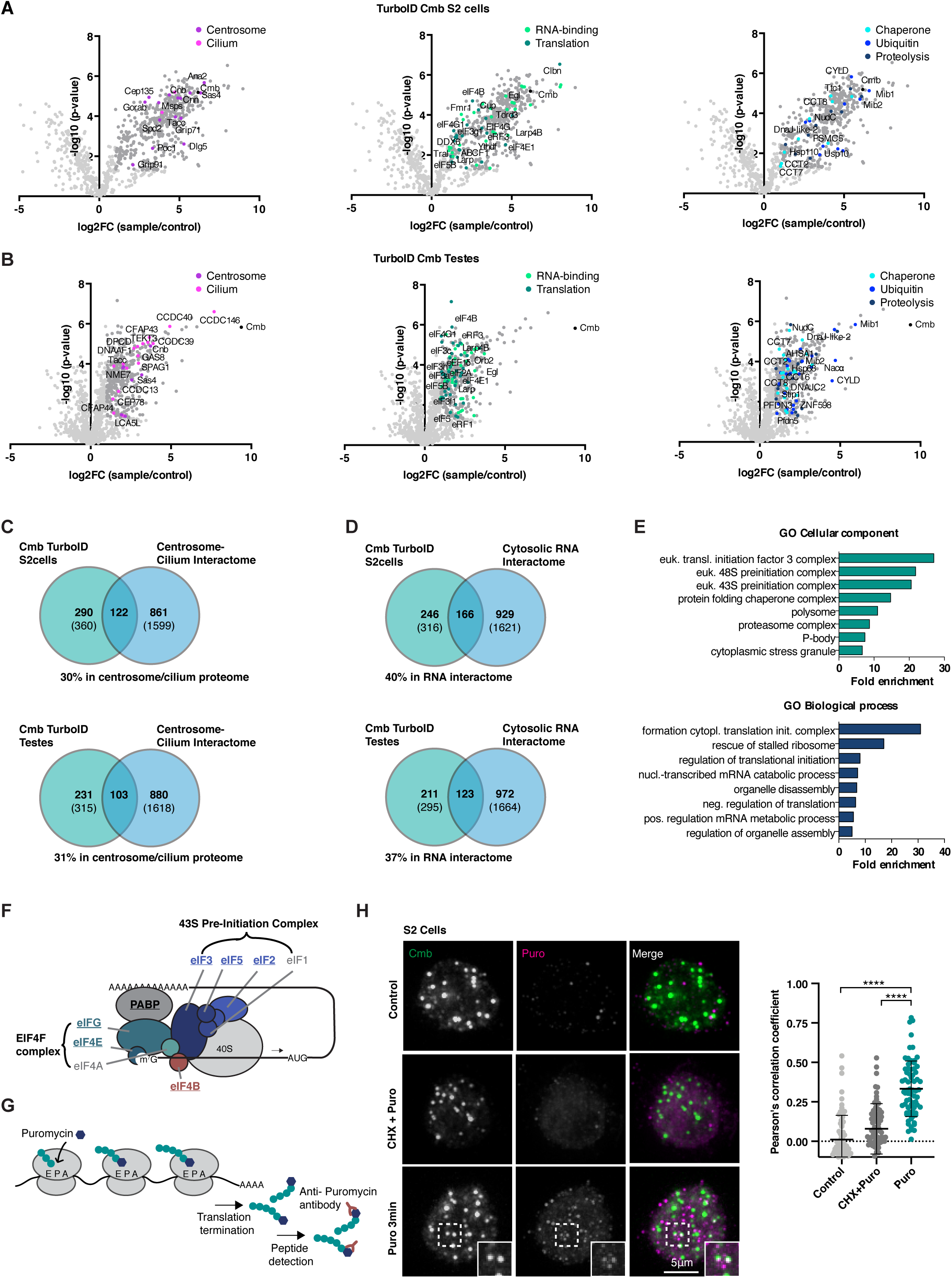
Cmb is associated with protein synthesis in *Drosophila*. (**A**, **B**) Direct TurboID of Cmb in S2 cells (cytosolic fraction, A) and indirect, GFP nanobody-targeted TurboID in fly testes (B) identifies centrosomal (magenta) and ciliary proteins (pink), RNA-binding proteins (light green) and proteins involved in translation (dark green), chaperone-mediated protein folding (light blue), ubiquitination (blue) and proteolysis (dark blue). Volcano plots of −log10 p-values against log2 fold change (sample/control). Significantly enriched proteins (Log2 enrichment >1, p-value <0.05) indicated in dark grey, with proteins of the above functional categories highlighted in color. See also Data S2B,D. (**C**) Venn diagrams showing overlap between Cmb S2 cell and testes TurboID interactomes and human centrosome/cilium proteome defined by (*3*). Comparison for those proteins conserved between human and flies. Numbers in parentheses are total number in each dataset. See also Data S2E. (**D**) Venn diagrams showing overlap between Cmb S2 cell and testes TurboID interactome and cytosolic RNA interactome defined by (*39*). Comparison for those proteins conserved between human and flies. Numbers in parentheses are total number in each dataset. See also Data S2E. (**E**) Gene Ontology (GO) enrichment analysis performed on human orthologs of the Cmb TurboID testes dataset. The top eight terms and their fold enrichments are shown for the GO categories ‘cellular component’ and ‘biological process’. (**F**) Schematic of eukaryotic translation initiation (*61*). The 43S-preinitiation complex is recruited to the mRNA by the EIF4F complex through interaction of EIF4G with eIF3. Components identified in the Cmb interactome are highlighted in bold. (**G**) Schematic of puromycin labeling assay. Puromycin mimics tyrosyl-tRNAs and binds the ribosomal acceptor site, blocking translation. Nascent peptide chains labeled with puromycin (puromycylated) are released into the cytoplasm and can be detected using antibodies against puromycin. (**H**) Puromycin labeling performed in *Drosophila* S2 cells. Puromycin labels Cmb foci in the cytoplasm after brief incubation with puromycin. No signal is detected in control cells or cells pre-treated with cycloheximide before addition of puromycin. Colocalization of Cmb and puromycin label quantified by Pearson’s correlation coefficient analysis. N=70 cells per condition. Kruskal-Wallis test followed by Dunn’s multiple comparison test used to assess statistical significance; ****P< 0.0001.

The identification of centrosomal and ciliary protein proximity interactors was to be expected. More surprising was the presence, in both flies and vertebrates, of mRNA-binding proteins, translation initiation factors and components of the protein quality control machinery (Figs 4A,B). Indeed, 37-40% of the Cmb interactome conserved in humans was also found in the cytosolic RNA interactome defined by (*39*), including components of the 43S pre-initiation complex, the most enriched GO term in our MS sample (Figs. 4D-F). Might satellites be associated with protein synthesis? To address this question we performed a puromycin incorporation assay (Fig. 4G). Puromycin, a structural analog of tyrosyl-tRNA, blocks translation by incorporating into the C-terminus of elongating polypeptide chains and triggering their release from ribosomes. Puromycin antibodies can be used to detect those polypeptide chains and, at short incubation times, identify sites of translation (*40*). Performing this assay in S2 cells revealed a remarkable degree of overlap of Cmb foci with puromycin label (Fig. 4H). Pre-treatment of cells with cycloheximide, which binds the ribosome and blocks eEF2-mediated translation elongation (*41*), abolished puromycin signal, confirming specificity. Cmb satellite foci, then, are sites of protein synthesis, at least in *Drosophila*. Could the same be true in vertebrates?

## Satellites are sites of translation of centrosomal and ciliary proteins in vertebrate cells

The puromycin incorporation assay revealed no clear cytoplasmic foci when performed in vertebrate cultured cells, potentially due to the rapid diffusion of nascent polypeptides away from the ribosome even at short incubation times (*42*). However, longer incubations with puromycin resulted in a loss of cytoplasmic signal for centrosomal and ciliary proteins, while centrosomal signal remained (Fig. 5A). The satellite population of these proteins is therefore newly synthesized. Based on the persistence of PCM1 signal, satellites themselves are unaffected, although they were now found dispersed throughout the cell (for an explanation of this phenomenon see below). Similarly unaffected were the ubiquitin ligase MIB1 and OFD1, a ciliopathy protein previously linked to protein synthesis and turnover (*43, 44*), which continued to colocalize with PCM1 (fig. S5A). This perturbation therefore appears to discriminate between centrosomal and ciliary clients and the machinery potentially involved in their synthesis.

**Fig. 5.**
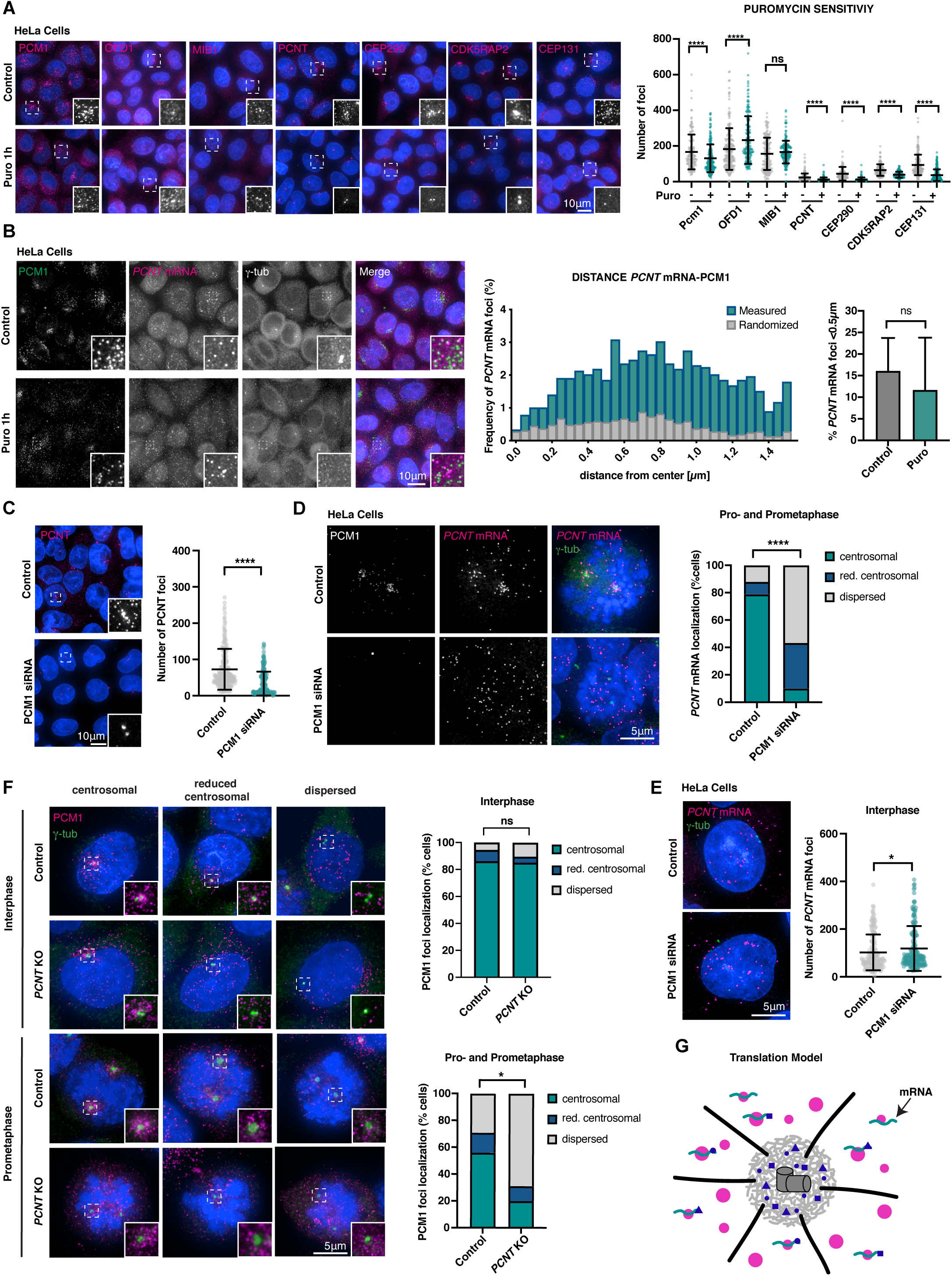
Centriolar satellites are sites of translation of centrosomal and ciliary proteins in vertebrate cells. (**A**) Immunofluorescence micrographs and quantitation of centriolar satellite protein signal in control and puromycin-treated Hela cells. Centrosomal and ciliary proteins PCNT, CEP290, CDK5RAP2 and CEP131 are significantly depleted of their cytoplasmic localization upon puromycin treatment, whereas their centrosome localization persists, suggesting the former represents newly-synthesized protein. In contrast, foci of PCM1, OFD1 and MIB1 remain, although now dispersed throughout the cytoplasm. >100 cells analyzed per condition. Mean +/-SD are displayed. Mann-Whitney test used to assess statistical significance; ****P< 0.0001. (**B**) Single-molecule fluorescence hybridization (smFISH) combined with immunofluorescence microscopy in control and puromycin-treated Hela cells. *PCNT* mRNA localizes in the vicinity of PCM1 independently of ongoing translation. To exclude random co-localization/proximity, distribution was compared with randomized controls. > 100 cells analyzed per condition. (**C**) Immunofluorescence micrographs and quantification of cytoplasmic PCNT signal in control and PCM1 siRNA-treated cells. Depletion of PCM1 leads to a significant decrease in number of PCNT foci (N=211 control cells and 171 PCM1 siRNA cells). Data analyzed using Mann-Whitney test; ****P< 0.0001. (**D**, **E**) smFISH combined with immunofluorescence microscopy in control cells and cells depleted of PCM1. PCM1 depletion impairs *PCNT* mRNA localization around the centrosome in prometaphase (D), reflecting a loss of co-translational targeting. However, the total number of *PCNT* mRNA foci assessed in interphase remains almost unchanged (E). N= 33 prometaphase-stage and 189 interphase cells control, 30 and 196 cells PCM1 siRNA. Mann Whitney test used to assess statistical significance; ****P< 0.0001, *P< 0.05. (**F**) Immunofluorescence micrographs of PCM1 in control and *PCNT* KO cells. Loss of PCNT does not impact PCM1 localization in interphase but compromises its concentration near centrosomes in pro-and prometaphase. N= 154 prometaphase and 348 interphase cells control, 161 and 355 cells *PCNT* KO. Localization groups compared using Mann-Whitney test, *P< 0.05. (**G**) Revised model of centriolar satellite function: centriolar satellites are sites of translation of centrosomal and ciliary proteins. Their concentration close to centrosomes in vertebrates is a byproduct of co-translational targeting of certain centrosomal/ciliary clients.

What the above results do not show is that translation of centrosomal and ciliary clients actually occurs at satellites, as opposed to satellites representing a waystation in the eventual translocation of newly synthesized protein to centrosomes. To distinguish between these possibilities, we performed fluorescence in situ hybridization to visualize the mRNA for PCNT, a PCM scaffolding component and satellite client protein (*6*). As seen in Figures 5B and S5B, *PCNT* mRNA localized in close proximity to both its cognate protein and PCM1, indicating that PCNT is translated in the immediate vicinity of satellites (see fig. S5C for a control of smFISH signal specificity). The same was true for CEP290, a satellite client and component of the ciliary transition zone (*45*) (not shown). The close proximity of *PCNT* mRNA to PCM1 remained in cells treated with puromycin and therefore devoid of cytoplasmic PCNT protein, indicating that PCM1 interacts with *PCNT* mRNA, not (or not only) protein (Fig. 5B). PCM1-containing satellites consequently act as sites where centrosomal and ciliary mRNAs concentrate and are potentially coordinately translated. The previously documented loss of satellite signal for centrosomal and ciliary clients following PCM1 depletion ((*6–8*), see Figs 5C, S5D) therefore reflects a failure to synthesize new proteins, not concentrate them in preparation for transit to centrosomes. Interestingly, putative components of the protein synthesis machinery are differentially affected by PCM1 depletion, with MIB1 dispersing throughout the cytoplasm, while OFD1, potentially in its role as a regulator of centriole elongation (*46*), continues to localize to centrosomes, while losing its cytoplasmic localization (fig. S5D). These results are consistent with PCM1 acting as the main scaffolding component of cytoplasmic translation factories specifically for centrosomal and ciliary proteins.

This left one key question: Why do satellites concentrate in the vicinity of centrosomes and the base of cilia in vertebrates? Clearly the fact that they do not in *Drosophila* indicates that satellite function in protein synthesis is independent of their localization. We were wondering whether this might be due to the previously reported phenomenon of co-translational targeting of certain centrosomal proteins including PCNT, which is recruited to centrosomes along with its mRNA at the onset of mitosis in a translation-, microtubule-and dynein-dependent manner (*47, 48*). *PCNT* mRNA highly concentrates at centrosomes in prophase/prometaphase, but then disperses in metaphase/anaphase, precisely the same pattern observed for PCM1 (*6, 48*). PCM1 depletion eliminates that enrichment without affecting mRNA abundance as assessed in interphase (Fig. 5D,E). In the original transport model of PCM1 function, this would reflect the role of satellites in moving nascent PCNT protein to centrosomes. In our alternative translation model, PCM1 would be a passive bystander in PCNT protein transport, while still engaged in synthesis of that protein. If so, PCNT depletion should affect PCM1 localization in mitosis, while in the original model PCM1 would be unaffected. The former is what we observed: PCM1 no longer concentrates at centrosomes in prophase/prometaphase of *PCNT* knockout cells (*49*), while its interphase localization is unaffected (Fig. 5F). The previously reported movement of satellites along microtubules (*9*) and their concentration at centrosomes in vertebrates is therefore a byproduct of co-translational targeting and incidental to their main function. PCNT is likely not the only satellite client that is co-translationally targeted. The dispersal of satellites following puromycin treatment in interphase cells (see above) indicates that their concentration near centrosomes at this stage is likewise a result of their function in translation of one or more such centrosomal clients, while such co-translational targeting does not seem to occur for their *Drosophila* counterparts.

## A revised model of centriolar satellite function

Since their discovery more than thirty years ago the specific functional contribution of centriolar satellites to centrosome and cilium biology has continued to elude researchers. Their seemingly dynamic localization pattern has inspired a variety of speculative hypotheses, of which the most prominent remains the transport model originally put forward in the early 2000s (*6*), according to which satellites are involved in microtubule-and dynein-dependent transport of centrosomal and ciliary cargo. This appeared to fit much of the data, but failed the most basic test: Satellites should move, yet, by and large, they do not (*14*). In our revised view of satellites as hubs of translation (Fig. 5G), movement occurs passively as a result of co-translational targeting of a subset of client proteins, but is neither driven by the core satellite machinery nor critical to their function. When viewed from this perspective, both their highly variable localization pattern in different cell types, cell cycle stages and environmental conditions (*6, 15, 16*) and their overall lack of movement begin to make sense. Depending on the levels of synthesis (and nature) of different client proteins satellites may concentrate in the vicinity of centrosomes, as they do in prophase/prometaphase, or be scattered throughout the cytoplasm. Thus, the reported dispersal of satellites in response to cellular stresses (*16*) is to be expected given the attendant shutdown of protein synthesis. The association of satellites with mRNAs encoding centrosomal and ciliary proteins is likely to also modulate the reported liquid-liquid phase separation behavior of satellites (*17, 18*). RNA is known to buffer phase separation behavior in the case of other membraneless organelles (*50*), and may act here to prevent the merger of satellites into large aggregates as occurs in certain PCM1 perturbations (*6, 34*). Such aggregation is known to perturb PCM1 function (*6*), and is likely to be disease relevant given the link of PCM1-containing satellites to ciliopathies and neurodegenerative disorders (*51, 52*).

The lack of any evident concentration of satellites in the vicinity of centrosomes or cilia in *Drosophila* shows that their specific intracellular localization is irrelevant for their function. In this respect protein synthesis at satellites differs from other reported instances of local translation, such as in the developing *Drosophila* embryo or in the axons and dendrites of vertebrate neurons, where it is thought to facilitate protein delivery to distant cellular locations or avoid inappropriate protein interactions en route (*53*). We suggest that the primary benefit of satellites is instead to facilitate the coordinate synthesis of members of the same multi-protein complexes for subsequent delivery to centrosomes and cilia. Such a role in refinement of centrosomal/ciliary protein expression is consistent with the rather minor yet highly pleiotropic consequences of PCM1 perturbation (*4, 5, 13*), another long-standing conundrum in the field. While satellites appear to be solely concerned with synthesis of centrosomal and ciliary proteins, no components of other cellular structures being reproducibly detected in the PCM1 or Cmb satellite proximity interactomes, it appears likely that coordinate synthesis also occurs for other non-membranous organelles. How selectivity is achieved in this case, as in other instances of local translation, remains unclear, but may involve primary sequence or secondary structure motifs (‘zip codes’, (*54, 55*)) present in centrosomal/ciliary mRNAs. Ribosome heterogeneity (*56*) may also play a role. We may not be returning to the extreme of ‘one gene-one ribosome-one protein’(*57*), but cellular protein synthesis appears to be much more organized than commonly appreciated. A better understanding of the role of proteins like PCM1 is clearly key to understanding how cells orchestrate the coordinate synthesis of the components that make up their various constituent structures.

## Acknowledgments

We thank members of the Dammermann and Campbell laboratories and Della David (Babraham Institute) for discussions; the Vienna *Drosophila* Resource Center (VDRC), the Bloomington *Drosophila* Stock Center (BDSC), Julius Brennecke and Jürgen Knoblich (IMBA), Andreas Jenny (Albert Einstein College of Medicine), Karen Oegema (UCSD), Norbert Perrimon (Harvard Medical School), and Jordan Raff (University of Oxford) for strains and reagents; Balazs Erdi of the IMBA fly house, Nicole Fellner of the VBCF Electron Microscopy facility, Weiqiang Chen and Markus Hartl of the Max Perutz Labs Mass Spectrometry facility and Josef Gotzmann and Thomas Peterbauer of the Max Perutz Labs BioOptics facility for technical assistance. Preliminary experiments for this project were performed by Astrid Steiner and Colette Emery.

## Author contributions

Conceptualization: CP, AD; Methodology: JGB (tools for image analysis); Investigation: CP (all other experiments), JD (*Drosophila* EM, initial behavioral and morphological analysis), CRK (initial BioID experiments), SR (orthology searches), MS (initial behavioral and morphological analysis), AB (PCNT KO immunofluorescence); Visualization: CP, AD; Funding acquisition: CP, JBG, AD; Project administration: AD; Supervision: AD; Writing – original draft: CP, AD; Writing – review & editing: all authors.

## Data and materials availability

Mass spectrometry data have been deposited to the ProteomeXchange Consortium and will be made freely available on publication. All other data are presented in the main text or supplementary materials. Requests for reagents should be directed to the corresponding author.

## Materials and Methods

### Materials used in this study

Key resources and tools used in this study are listed in Tables S1-S7:

Table S1. Fly strains

Table S2. Cell lines

Table S3. Plasmid constructs

Table S4. Antibodies

Table S5. Chemicals, enzymes and other reagents

Table S6. Oligonucleotides

Table S7. Software

### *Drosophila* melanogaster stocks and husbandry

Cmb mutants and flies overexpressing Cmb-RA under GAL4/UAS control were a gift from Andreas Jenny, flies expressing RFP-Cnn (*62*), H2A-RFP (*63*) and Asl-GFP (*64*) from Jordan Raff, Bam-GAL4 (*65*) and Pnr-GAL4 (*66*) driver lines from Helen White-Cooper and Jürgen Knoblich, respectively. Flies expressing GFP nanobody-TurboID under the *eggless* promoter (derived from an NLS-containing construct generated by the lab of Julius Brennecke, (*67*)) were generated by integration into the attP40 landing site on Chromosome 2 by the IMBA Fly Facility. All other fly strains were obtained from the *Drosophila* Stock Center (VDRC) or Bloomington *Drosophila* Stock Center (BDSC) or generated by genetic crosses as detailed in Table S1. Further information on strains is available on FlyBase (http://flybase.org), as well as on the websites of the two stock centers (BDSC, https://bdsc.indiana.edu, VDRC, https://stockcenter.vdrc.at/control/main). Flies were maintained on standard media at 25°C. W^1118^ flies were used as controls for all experiments. For RNAi, UAS-hairpin RNAi males and GAL4 driver line females were crossed in standard vials and allowed to mate for 4 days at 25°C, then shifted to 29°C to induce GAL4 expression.

### Insect and vertebrate cell culture

*Drosophila* Schneider S2 cells were obtained from Life Technologies and cultured in Schneider’s medium containing 10% FBS, penicillin (50 units/ml), and streptomycin (50µg/ml). Cells were kept at 25°C at atmospheric CO_2_ and passaged every 3-4 days. For proximity interaction analysis of Sas4 by BioID in S2 cells, a construct expressing myc-BirA*(R118G) under the copper-inducible metallothionein promoter (pMt-myc-BirA*) was generated by inserting the myc-BirA* coding sequence (derived from a plasmid obtained from Kyle Roux) into the pMt-V5-6xHisB vector backbone (Invitrogen). The coding sequence of Sas4 was then inserted into that construct by PCR and restriction cloning. For Cmb TurboID, the coding sequence of Cmb-RA was cloned into the pDONR Zeo entry vector (Invitrogen) and then transferred into the pMT-TurboID-V5 vector (a gift from Norbert Perrimon) by Gateway cloning. To monitor Cmb dynamics in S2 cells, the coding sequence of Cmb-RA was cloned into pMt-V5-6xHisB by Gibson cloning. Clonal cell lines expressing GFP/biotin ligase fusions were generated using TransIT transfection reagent (Mirus) by co-transfection of 0.6µg/ml expression plasmid with 0.06µg/ml pCoBLAST selection plasmid (Invitrogen). Cells were grown for 3-5 days before selection with 25µg/ml blasticidin. GFP/biotin ligase expression was induced by addition of 500/100µM of CuSO_4_ for 24h before imaging/harvesting.

Puromycin labeling was carried out as described in (*68*). Cells were treated with 10µg/ml puromycin for different incubation times (30sec, 1min, 3min, 5min, 10min). Control cells were treated with DMSO for the same period of time or pre-treated with 10µg/ml Cycloheximide (CHX) for 30min prior to addition of puromycin. After treatment cells were immediately fixed in −20°C methanol and processed for immunofluorescence as described below.

HeLa CCL-2 cervical carcinoma cells and their derivative *PCNT* knockout cell line ODCL0025 (*49*) were a gift from Karen Oegema and cultured in DMEM containing 10% FBS, 25mM Hepes (Gibco) and non-essential amino acids (Gibco). Human osteosarcoma U2-OS HTB-96 cells are from the American Type Culture Collection (ATCC) and were cultured in Dulbecco’s Modification of Eagles Medium (DMEM) containing 10% FBS, penicillin (100 units/ml), and streptomycin (0.1mg/ml). Cells were grown at 37°C in 5% CO_2_ and passaged every 2-3 days. All cell lines were regularly tested for mycoplasma contamination.

RNAi-mediated depletion of PCM1 was performed by siRNA using a mixture of two oligonucleotides ((*6*), see Table S6), purchased from Sigma-Aldrich and introduced into cells by Lipofectamine RNAiMAX-mediated transfection following manufacturer’s instructions. Phenotypes were assessed 48 hours after transfection.

### Identification of orthologs of centrosomal/ciliary genes across opisthokonts

PCM1 orthologs, as well as orthologs of centrosomal/ciliary genes across different phyla were identified by reciprocal BLAST search (BLAST+ 2.13.0, (*69*)) performed locally using the human protein as the starting point, with bidirectional best match at an E-value threshold cutoff of 0.1 as a simple but robust (*70*) method to infer orthology. Where direct comparisons failed to identify a clear ortholog, indirect searches were performed using less divergent related species as intermediates, as well as by applying hidden Markov models with HMMER (*71*) using alignments of known representatives, constructed with MAFFT (*72*). Conserved motifs within PCM1 were identified by Meme (https://meme-suite.org/meme/) and sequences aligned using MUSCLE within Jalview (https://www.jalview.org). The presence of centrosomes and cilia within fungi, Nematomorpha and Platyhelminthes was inferred from the conservation of core centriolar (STIL/Ana2, SASS6/Sas6, CENPJ/Sas4, CEP135/Bld10), centrosomal (CDK5RAP2/Cnn, CEP192/Spd2) and ciliary (transition zone, IFT and BBS components, inner and outer dynein arm components, dynein assembly factors, nexins, N-DRC, radial spoke and central apparatus components, (*28*)) proteins, as well as literature reports (*21, 58*).

### *Drosophila* behavioral assays

To examine fly coordination, 10 three-day-old adult males were collected in 8cm graduated, flat bottom tubes. Flies were left for 30min to recover from the anesthetic and acclimatize, then banged to the bottom of the tube and filmed climbing back upward. Videos were analyzed to establish the time at which all flies had crossed the half-way mark. Experiment was repeated 3 times per condition.

To assess male fertility, single three-day-old males were crossed with four control virgin females in standard culture vials. After mating for 1 hour, males were removed and females kept in the vial for an additional 24 hours and the number of offspring counted prior to hatching. Experiment was repeated 3 times with 10 males per condition.

To assess embryonic viability, two virgin females and one male were put into cages with apple juice plates and allowed to lay eggs for 24 hours at 25°C. Agar plates were transferred into a humid chamber and the number of hatched and unhatched embryos determined after 24 and 48 hours.

### *Drosophila* immunofluorescence staining

Rabbit polyclonal antibodies against the N-terminus (amino acids 1-150) of *Drosophila* Cmb were raised using a GST fusion as antigen and purified over the untagged cleaved antigen as described (*73*). The following primary antibodies were used for immunofluorescence in *Drosophila*: mouse anti-puromycin (clone 12D10, Merck Millipore) at 1:200, rabbit anti-Cmb antibody (this study), 1:300, mouse anti-NompC (*74*), rabbit anti-Ana1 (*62*), rabbit anti-Cep97 (*27*), rabbit anti-Sas4 (*30*), all at 1:500, rabbit anti-Cnn (*75*), 1:600, rabbit anti-CP110 (*76*), 1:700, rabbit anti-Spd2 (*77*), 1:800, rabbit anti-Ana2 (*77*), 1:900, mouse anti-γ-tubulin (GTU88; Sigma-Aldrich), 1:1000. Secondary antibodies (Jackson ImmunoResearch) were used at 1:100.

For immunofluorescence staining in S2 cells, cells seeded on Concavalin A-coated coverslips were fixed in 4% formaldehyde for 15min at room temperature (or −20°C methanol for 20min in the case of puromycin labeling), permeabilized with PBS 0.2% Tween-20 for 15min and blocked 20min in 5% BSA in PBS 0.1% Triton-X100. Cells were incubated with primary antibodies in blocking solution for 1 hour, washed 3x 5min with PBS 0.1% Triton-X100 and incubated with secondary antibodies in blocking solution for 1 hour. Following staining with 1µg/ml Hoechst in PBS for 5min cells were washed 3x 5min with PBS 0.1% Triton-X100 and once with PBS before being mounted in Vectashield.

For immunofluorescence staining in testes, adult males were dissected in ice-cold PBS, the testes transferred to a microscope slide, covered with a coverslip and flash-frozen in liquid nitrogen. After recovering slides and removing the coverslip, samples were incubated for 5min in −20°C methanol and 1-2min in −20°C acetone. Samples were then rinsed with PBS, rehydrated in PBS 1% Triton X-100 for 10min and washed 2x with PBS for 5min, followed by blocking with 1% BSA in PBS for 45min. Primary antibodies in blocking solution were then added and samples incubated overnight at 4°C. The next day, samples were washed 2x with PBS for 5min before incubation with secondary antibodies in blocking solution for 1 hour. Finally, after 2x washes with PBS for 5min, samples were stained with 1µg/ml Hoechst in PBS for 5min and mounted in Vectashield.

For immunofluorescence staining of chordotonal neurons, leg chordotonal organs were dissected out of 36 hour old male pupae, placed between a microscope slide and coverslip and gently pressed to remove them from the cuticle. Fixation and staining were performed as described for testes above.

For immunofluorescence staining of neuroblasts, brains from third-instar larvae were dissected in PBS and placed in 4% paraformaldehyde in PBS for 20min, treated with 45% acetic acid for 15s, followed by 60% acetic acid for 3min, then covered with a coverslip and squashed between coverslip and slide before flash-freezing in liquid nitrogen. After recovering slides and removing the coverslip, samples were fixed in methanol for 5-8min at −20°C, washed 4x 15min with PBS 0.1% Triton-X100 and incubated in primary antibody in PBS 0.1% Triton-X100 overnight at 4°C. The next day, brains were washed 3x 5min with PBS 0.1% Triton-X100 and incubated with secondary antibody overnight at room temperature for 3-4 hours. Finally, samples were washed 3x 15min with PBS 0.1% Triton-X100, stained with 1µg/ml Hoechst in PBS for 5min and mounted in Vectashield.

For embryo immunofluorescence, flies were put into cages on apple-juice plates with yeast-paste 3 days before embryo collection. Early embryos (0-2 hours) were collected from plates, washed and quickly treated with 5% bleach to dechorionate them and fixed in heptane/4% paraformaldehyde in PBS for 2min. Heptane was replaced by methanol and samples incubated for 1min. Embryos were rehydrated with PBS 0.1% Triton X-100 for 15min rotating, blocked 2x with 5% BSA in PBS for 30min before incubation with primary antibody in blocking buffer overnight at 4°C. The next day, embryos were washed 3x with PBS 0.1% Triton X-100 for 20min and incubated on the rotator with secondary antibodies in blocking buffer for 1 hour. Finally, embryos were washed in PBST, stained with 1µg/ml Hoechst in PBS for 5min, washed again and mounted in Vectashield.

### *Drosophila* wholemount testes staining

For Hoechst/Phalloidin wholemount testes staining, adult males were dissected in PBS and testes transferred to Nunc MicroWell MiniTrays containing PBS and fixed in 4% paraformaldehyde in PBS for 25min. Testes were washed 3x with PBS 0.1% Triton-X100 for 5min and stained with 5µg/ml Hoechst and 1:50 Phalloidin Alexa Fluor 568 in PBS overnight at 4°C. The next day, testes were washed 3x with PBS, carefully separated from the rest of the abdomen and placed in a drop of Vectashield on a multi well slide and covered with a coverslip.

### Immunofluorescence staining in vertebrate cells

The following primary antibodies were used for immunofluorescence in vertebrate cells: rabbit anti-MIB1 (Sigma-Aldrich), 1:100, rabbit anti-AZI1 (Abcam), 1:250, mouse anti-hsPCM1 (G-6, Santa Cruz), rabbit anti-PCM1 (*6*), both 1:300, rabbit anti-CDK5RAP2 (Merck Millipore), rabbit anti-CEP290 (Novus Biologicals), rabbit anti-PCNT (Abcam), all 1:400, rabbit anti-OFD1 (Sigma-Aldrich), 1:500, mouse anti-γ-tubulin (GTU88; Sigma-Aldrich), 1:1000. Secondary antibodies (Jackson ImmunoResearch) were used at 1:100.

For immunostaining in vertebrate cells, cells were fixed in −20°C methanol for 20min, then washed and rehydrated in PBS 2x 5min. Permeabilization was done in PBS + 0.1% Tween-20 for 5min, followed by blocking in 0.5% BSA in PBS-Tween-20 for 5min. Cells were subsequently incubated with primary antibodies in blocking buffer for 40min, washed 2x 5min with PBS and once with PBS-Tween-20. Cells were then incubated with secondary antibodies for 20min and stained with 1µg/ml Hoechst in PBS for 5min. After 2 further washes with PBS (5min each) and once with PBS-Tween-20, coverslips were mounted on slides with Vectashield. For immunofluorescence experiments including the mouse anti-hsPCM1 (G-6, Santa Cruz) antibody, we followed the protocol described in (*78*).

### Sequential single molecule fluorescence hybridization and immunofluorescence staining in vertebrate cells

Fluorescence in situ hybridization probes against the coding sequence of *PCNT* mRNA (transcript variant 1, Accession: NM_006031) were designed using Stellaris Probe Designer (www.biosearchtech.com/stellarisdesigner) and ordered conjugated with Quasar 570. Sequences can be found in Table S6. Probes were resuspended in TE buffer at 25mM. The protocol for sequential smFISH-IF was adapted from (*79*) and (*80*). Cells were fixed in 4% paraformaldehyde for 15min, washed 3x 10min with PBS 0.05% Triton-X100, then incubated in warmed pre-hybridisation buffer/FISH wash buffer (10% formamide, 2x SSC) at 37°C for 20min before incubation in hybridization buffer (10% formamide, 10mg/ml sheared salmon sperm DNA, 10% dextran sulfate, 10mM Vanadyl ribonucleoside complexes solution) containing 300nM smFISH probe overnight at 37°C. The next day cells were permeabilized and blocked with 1% BSA in 2x SSC 1% Triton-X100 for 2 hours (solution exchanged twice during incubation). Cells were then washed 2x with 2x SSC and incubated with primary antibody in 2x SSC and 1% Triton-X100 for 1 hour, washed 3x with IF wash buffer (2x SSC, 0.5% Triton-X100 in ddH2O), and incubated with secondary antibody solution in 2x SSC for 2-4 hours at room temperature. Finally, cells were washed 3x 10min with IF wash buffer, stained with 1µg/ml Hoechst in PBS for 5min and mounted in Vectashield.

### Fixed imaging

For imaging of fixed samples (except wholemount testes staining), 0.2µm 3D widefield datasets were acquired using either a 60x 1.42NA UPLanX Apochromat or a 100x 1.4NA UplanS Apochromat lens on a DeltaVision Ultra Epifluorescence Microscope equipped with a 4-Megapixel sCMOS camera and 7-Color SSI module, computationally deconvolved using SoftWorx and imported into Fiji for post-acquisition processing. Exposure settings were held constant across conditions.

For wholemount testes staining, samples were imaged on a Zeiss LSM 700 confocal laser scanning microscope equipped with 40× 1.3NA Plan Apochromat oil immersion objective. Image stacks of approximately 20 z-planes were acquired at 4 µm increments for each of the six testes. Maximum projections from 4 z-stacks were stitched together using ZEN software (Zeiss) to cover the whole testis.

### Live cell imaging

For live cell imaging of *Drosophila* syncytial-stage embryos, 0-2 hour embryos were collected on apple juice agar plates, dechorionated on double-sided tape, immobilized on glass-bottom dishes coated with Scotch tape adhesive dissolved in heptane and mounted with Voltalef oil. Embryos were imaged on a Yokogawa CSU-X1 spinning disk confocal mounted on a Zeiss Axio Observer Z1 inverted microscope equipped with a 63x 1.4NA Plan Apochromat lens, 100mW 488nm and 200mW 561nm solid state lasers and a Hamamatsu ImageEM X2 EM-CCD camera and controlled by VisiView software (Visitron Systems). 14×0.75µm GFP/mCherry z-series were acquired at 15s intervals, with manual focus adjustment between intervals if needed, using low laser illumination to minimize photobleaching. Image stacks were imported into Fiji for post-acquisition processing.

For live cell imaging in S2 cells, GFP expression was induced with CuSO4 24hours before imaging. On the day of imaging, cells were plated on glass bottom dishes and allowed to settle for at least 30min. Imaging was performed on a Yokogawa CSU X1 spinning disk confocal mounted on a Zeiss Axio Observer Z1 inverted microscope equipped with a 100x 1.3NA EC Plan-Neofluar lens, 100mW 488nm solid-state laser and an Evolve EM512 EMCCD camera (Photometrics) and controlled by VisiView software (Visitron Systems). 0.5µm GFP z-series were acquired at 5s intervals, using low laser illumination to minimize photobleaching. Image stacks were imported into Fiji for post-acquisition processing.

For live cell imaging of sperm motility, seminal vesicles of 3-day old males were dissected in PBS and immediately transferred into a drop of Schneider medium on a microscope slide. The seminal vesicle was pierced with a sharp tungsten needle, mineral oil used to encircle the drop, a coverslip placed on top and sperm motility imaged immediately using dark field settings on a Zeiss Axio Imager Z2 microscope equipped with a 63x 1.4NA Plan-Apochromat lens at 12 frames/s. Image sequences were imported into Fiji for post-acquisition processing.

### Quantification of centrosome intensity in the early embryo

Analysis of centrosome intensity in the early embryo was done as described in (*81*) with modifications described hereafter. Maximum intensity projections of z-stacks were generated and image sequences bleach-corrected using the ImageJ plugin ‘Bleach correction’ (*82*) applying the exponential fitting method. Centrosome intensity (RFP:Cnn channel) was analyzed at nuclear envelope breakdown (NEBD). A fixed-sized ROI (4.88µm x 4.88µm) was placed manually around the center of each centrosome and the mean intensity of this centrosome ROI computed using Fiji. For background quantifications, we analyzed mean intensities within the same sized ROI placed in three cytoplasmic regions close to the centrosome, calculated the average and subtracted this value from the centrosome mean intensity. For each embryo, all centrosomes that were fully captured within the z-stack were used for quantification and included in the statistical test. The final average mean intensity per embryo was calculated and plotted in GraphPad Prism.

### Colocalization analysis

To assess colocalization of Cmb/PCM1 with proteins of interest the Fiji colocalization Plugin JaCoP (*83*) was used for both image channels. A cytoplasmic region containing satellite foci away from the nucleus of size 3.5µm x 3.5µm was defined in Fiji for both channels. To assess colocalization at the centrosome, the ROI for both channels was positioned such that the centrosome was located not in the center but at one extreme of the ROI. The 3D z-stack was cropped and segmented using the built-in threshold option. To exclude random colocalization, one of the channels was rotated by 90°. The Pearson’s correlation coefficient was calculated and used as an indicator of the degree of overlap. Any difference between experimental and randomized condition was assessed by statistical testing.

### Quantification of number of foci

For quantification of number of satellite or mRNA foci, maximum projections were generated in Fiji. Pixel-based segmentation of foci was carried out using Ilastik with the simple segmentation method. A custom-made Fiji macro was used to convert the background and signal image classes into 0 and 1s, respectively. The segmented images were analyzed in Matlab with a custom-made script. In brief, the maximum intensity projections of DAPI, centrosome and foci channels were combined into a single image, the user given the opportunity to adjust brightness and contrast to make the outline of the cells visible and the outline of individual cells manually delineated using the CROIEditor.m function (https://github.com/aether-lab/prana/blob/master/CROIEditor.m). Segmented images were post-processed using the watershed algorithm and the area and intensity of foci quantified using regionprops. The cell background intensity was calculated, results for intensities, number of foci per cell and foci size imported into a.csv table and analyzed in Excel.

### Proximity analysis for mRNA and protein

To assess proximity of mRNA and protein the object-based Fiji plugin DiAna (*84*) was used. While DiAna is capable of proximity measurements in 3D, a 2D measurement was chosen in order to take advantage of the ‘shuffle’ function to test for non-random proximity. Deconvolved, single-plane images for the two respective channels (mRNA and protein) were loaded into Fiji and segmented using global thresholding and Gaussian filtering implemented in the plugin. Distance to the first nearest object (center-center distance) was calculated. To assess statistical significance, distances between pairs of proximal particles was compared to their counterparts in randomized images. For this purpose, a mask was applied using Fiji ‘create mask’ to manually outline each cell (excluding the nucleus, mitotic and apoptotic cells). In randomized conditions, the ‘shuffle’ function was used to randomly distribute objects within the mask and distances calculated as described before.

### Quantification of satellite signal around centrosomes

To determine the subcellular distribution of PCM1 and Cmb relative to the centrosome, radial profiles were quantified. First, maximum intensity projections were created, thresholded and the center of mass of each centrosome based on γ-tubulin signal identified using the ‘analyze particles’ function in Fiji. A circular ROI was placed around this point and the plugin Radial Profile Plot (https://imagej.nih.gov/ij/plugins/radial-profile.html) used to obtain a radial intensity profile of satellite foci centered around each centrosome. Concentric rings and corresponding integrated intensities were quantified within 5µm of the centrosome. For background quantifications, three smaller sized circular ROIs in distant cytoplasmic regions without foci were used to subtract from the mean intensity value. Intensities were then normalized to the central ring, all intensities were combined, averaged and mirrored to plot a single symmetric radial profile around the centrosome.

### Single particle tracking and analysis of satellite dynamics

Time-lapse recordings of Cmb foci were first exported and processed using Fiji to create maximum intensity projections. Segmentation of Cmb foci was performed using Ilastik (v1.4.0-OX) (*85*) and the simple segmentation method (*86*). Particles exceeding 700nm in diameter or exceeding a mean fluorescence signal intensity two-fold higher than average were excluded from analysis to avoid artefacts of overexpression. Single particle tracking of Cmb-GFP satellites in S2 cells was performed using Trackmate (v7.11.1) (*87*) with the following settings: Ilastik detector, simple linear assignment problem (LAP) tracker, maximum linking distance of 5µm, maximum gap-closing distance of 5µm and maximum gap-closing of 2 frames. The resulting spots and tracks features were exported and further analyzed in Excel. Instantaneous speed, average speed and mean squared displacement were computed from coordinates of satellites from trajectory analysis. Tracks were further characterized using TraJClassifier (v0.8.1) (*88*), a Fiji plugin that classifies trajectories into normal diffusion, subdiffusion, confined diffusion and directed/active motion by a random forest approach. Parameters used were as follows: minimum track length of 11 frames, window size (positions) of 10, segment length of 10, resample rate and pixel size 0 and frame rate of 0.2 frames per second.

### *Drosophila* transmission electron microscopy

Transmission electron microscopy was carried out as previously described in (*27*). For EM of chordotonal organs, legs from 36hour old pupae were cut off with microscissors and fixed in 2% glutaraldehyde and 2% paraformaldehyde in 0.1mol/l sodium phosphate buffer (pH 7.2) in a desiccator for 2 hours at room temperature and then overnight on a rotator at 4°C. The next day, legs were rinsed with sodium phosphate buffer, post-fixed in 2% osmium tetroxide in buffer on ice, dehydrated in graded series of acetone on ice and subsequently embedded in Agar 100 resin. 70nm sections were cut and post-stained with 2% uranyl-acetate and Reynolds lead citrate. Sections were examined on a Morgagni 268D microscope (FEI) operated at 80kV. Images were acquired with a 11 megapixel Morada CCD camera (Olympus-SIS).

For EM of testes, late pupal testes from third instar larvae were dissected in PBS and fixed using 2.5% glutaraldehyde in 0.1mol/l sodium phosphate buffer, pH7.2 for 1 hour at room temperature. Samples were then rinsed with sodium phosphate buffer, post-fixed in 2% osmium tetroxide in ddH2O on ice, dehydrated in a graded series of acetone and embedded in Agar 100 resin. 70nm sections were then cut, processed and imaged as described above.

### BioID and TurboID in *Drosophila* S2 cells

BioID and TurboID pulldown was performed as detailed in (*89*), with minor modifications. Cells expressing pMt-myc-BirA*, pMt-Sas4-myc-BirA*, pMt-V5-TurboID-GW, or pMt-V5-TurboID-Cmb were incubated overnight with 50μm biotin, washed 3x with PBS and lysed in ELB+ buffer (150mM NaCl, 50mM Hepes pH7.5, 5mM EDTA, 0.3% NP-40, 6% Glycerol) supplemented with Roche cOmplete Mini EDTA-free Protease-Inhibitor-Cocktail (1 tablet/10ml lysis buffer), 1mM PMSF and 1mM Benzamidine. Lysates were clarified by brief centrifugation at 200g for 3min at 4°C in a benchtop centrifuge before pelleting insoluble cellular material including centrosomes by centrifugation at 20000g for 30min at 4°C. To perform TurboID on the detergent-soluble cytoplasmic fraction, supernatant was used directly for pulldown. For the detergent-insoluble cytoskeletal fraction, pellets were resuspended in 2% SDS, 1% beta-mercaptoethanol in PBS and boiled for 30min at 95°C with intermittent vortexing. Subsequently, the SDS concentration was reduced to 0.2% by dilution with PBS and Triton-X-100 added to a final concentration of 2%. Samples were then sonicated by tip sonication (3x 30s pulses using a Bandelin Sonopuls GM70 sonicator at 60% continuous output, with brief cooling on ice between pulses), the SDS concentration reduced to 0.1% SDS with PBS and samples sonicated once more for 30s at 60% output before centrifuging at 20000g for 30min at 4°C and recovering the supernatant. Pierce Streptavidin-coated magnetic beads were equilibrated with PBS containing 0.1% SDS before incubating with the detergent-soluble cytoplasmic fraction or solubilized cytoskeletal fraction on a rotator overnight at 4 °C. For Cmb TurboID samples, beads were further acetylated to reduce background (*90*) by incubation with a mixture of 190μl 50mM HEPES-NaOH pH7.8 containing 0.2% Tween-20 and 10μl 100mM Pierce Sulfo-NHS-Acetate in for 1 hour at room temperature for 1h prior to incubation with lysate. Following the overnight incubation, unbound lysate was removed and beads washed for 8min each on a rotator at room temperature, first with 2% SDS in ddH_2_O, then 0.1% deoxycholic acid, 1% Triton X-100, 1mM EDTA, 500mM NaCl and 50mM HEPES pH7.5, and finally 0.5% deoxycholic acid, 0.5% NP-40, 1mM EDTA, 500mM LiCl and 10mM Tris pH7.5. Beads were then washed 5x for 3min with 50mM Tris pH7.4 before being sent for on-bead protein digestion and mass spectrometry analysis.

### Nanobody TurboID *Drosophila* testes

Lysates were prepared as previously described for *C. elegans* in (*36*). Briefly, adult flies were washed 3x with PBS, recovering flies by centrifugation for 3min at 300g, quickly ground in RIPA buffer (1% Triton X-100, 1mM EDTA, 0.5% sodium deoxycholate, 0.1% SDS, 150mM NaCl, 50mM Tris-HCl pH7.4) supplemented with Roche cOmplete Mini EDTA-free Protease-Inhibitor-Cocktail (1 tablet/10ml lysis buffer), 1mM PMSF and 1mM Benzamidine using a mortar and pestle and drop-frozen in liquid nitrogen. Frozen fly ‘popcorn’ was further ground by cryogenic milling using a SPEX 6875 cryogenic mill (5 cycles, 1min pre-cool, 2min run-time, 1min cool-time; 12 cps). Lysates were thawed and SDS and DTT added to a final concentration of 1% and 10mM, respectively, boiled at 95°C for 5min and sonicated by tip sonication (2x 1min at 20% continuous output, with brief cooling on ice between pulses). Urea solution (8M urea, 1% SDS, 50mM Tris-Cl, 150mM NaCl) was added to a final concentration of 2M urea and lysates centrifuged at 100000g for 45 min at 22°C. Lysates were desalted over Zeba spin desalting columns (7K MWCO, Thermofisher) to remove free biotin and incubated with streptavidin magnetic beads by incubating on a rotator overnight. Following incubation, beads were washed twice with 150mM NaCl, 1mM EDTA, 2% SDS, 50mM Tris-HCl, pH7.4, once with 1x TBS buffer pH 7.4, twice with 1M KCl, 1mM EDTA, 50mM Tris-HCl, 0.1% Tween-20, pH7.4, twice with 0.1M Na_2_CO_3_, 0.1% Tween-20, pH 11.5, twice with 2M urea, 10mM Tris-HCl, 0.1% Tween-20 pH 8.0 and finally 5x with 1x TBS buffer. Beads were finally resuspended in 1X TBS buffer and sent for on-bead protein digestion and mass spectrometry analysis.

### Sample preparation for mass spectrometry analysis

Beads were resuspended in 50µl 1M urea and 50mM ammonium bicarbonate. Disulfide bonds were reduced with 2µl of 250mM DTT for 30min at room temperature before adding 2µl 500mM iodoacetamide and incubating for 30min at room temperature in the dark. Remaining iodoacetamide was quenched with 1µl of 250mM DTT for 10min. Proteins were digested with 150ng LysC (mass spectrometry grade, FUJIFILM Wako chemicals) in 1.5µl 50 mM ammonium bicarbonate at 25°C overnight. The supernatant without beads was digested with 150ng trypsin (Trypsin Gold, Promega) in 1.5µl 50mM ammonium bicarbonate followed by incubation at 37°C for 5 hours. The digest was stopped by the addition of trifluoroacetic acid to a final concentration of 0.5%, and the peptides desalted using C18 Stagetips (*91*).

### Liquid chromatography separation coupled to mass spectrometry

Peptides were separated on an Ultimate 3000 RSLC nano-flow chromatography system (Thermo-Fisher), using a pre-column for sample loading (Acclaim PepMap C18, 2cm × 0.1mm, 5μm, Thermo-Fisher), and a C18 analytical column (Acclaim PepMap C18, 50cm × 0.75mm, 2μm, Thermo-Fisher), applying a segmented linear gradient from 2% to 35% and finally 80% solvent B (80% acetonitrile, 0.1% formic acid; solvent A 0.1% formic acid) at a flow rate of 230nl/min over 120min. The peptides eluted from the nano-LC were analyzed by mass spectrometry as described below.

For Sas4 S2 cell BioID, a Q Exactive HF Orbitrap mass spectrometer (Thermo Fisher) coupled to the column with a nano-spray ion-source using coated emitter tips (PepSep, MSWil), was used with the following settings: The mass spectrometer was operated in data-dependent acquisition mode (DDA), survey scans were obtained in a mass range of 380-1650m/z with lock mass activated, at a resolution of 120k at 200m/z and an AGC target value of 3E6. The 10 most intense ions were selected with an isolation width of 2.0m/z without offset, fragmented in the HCD cell at 27% collision energy and the spectra recorded for max. 250ms at a target value of 1E5 and a resolution of 30k. Peptides with a charge of +2 to +6 were included for fragmentation, the peptide match and the exclude isotopes features enabled, and selected precursors were dynamically excluded from repeated sampling for 30s.

For Cmb S2 cell TurboID, a Q Exactive HF-X Orbitrap mass spectrometer (Thermo Fisher) coupled to the column with a nano-spray ion-source using coated emitter tips (PepSep, MSWil), was used with the following settings: The mass spectrometer was operated in data-dependent acquisition mode (DDA), survey scans were obtained in a mass range of 375-1500m/z with lock mass activated, at a resolution of 120k at 200m/z and an AGC target value of 3E6. The 8 most intense ions were selected with an isolation width of 1.6m/z with offset 0.2m/z, fragmented in the HCD cell at 28% collision energy and the spectra recorded for max. 250ms at a target value of 1E5 and a resolution of 30k. Peptides with a charge of +2 to +6 were included for fragmentation, the peptide match and the exclude isotopes features enabled, and selected precursors were dynamically excluded from repeated sampling for 30s.

For Cmb testes TurboID, an Exploris 480 Orbitrap mass spectrometer (Thermo Fisher) coupled to the column with a FAIMS pro ion-source (Thermo-Fisher) using coated emitter tips (PepSep, MSWil), was used with the following settings: The mass spectrometer was operated in DDA mode with two FAIMS compensation voltages (CV) set to −45 or −60 and 1.5s cycle time per CV. The survey scans were obtained in a mass range of 350-1500m/z, at a resolution of 60k at 200m/z and a normalized AGC target at 100%. The most intense ions were selected with an isolation width of 1.2m/z, fragmented in the HCD cell at 28% collision energy and the spectra recorded for max. 100ms at a normalized AGC target of 100% and a resolution of 15k. Peptides with a charge of +2 to +6 were included for fragmentation, the peptide match feature was set to preferred, the exclude isotope feature was enabled, and selected precursors were dynamically excluded from repeated sampling for 45s.

### Mass spectrometry data analysis

The RAW MS data were analyzed with FragPipe (20.0), using MSFragger (3.8) (*92*), IonQuant (1.9.8) (*93*), and Philosopher (5.0.0) (*94*). The default FragPipe workflow for label free quantification (LFQ-MBR) was used, except “Normalize intensity across runs” was turned off. For Sas4 BioID, the “MBR top runs” parameter was set to 1 to address batch measurements of sample replicates. Cleavage specificity was set to Trypsin/P, with two missed cleavages allowed. The protein FDR was set to 1%. A mass of 57.02146 (carbamidomethyl) was used as fixed cysteine modification; methionine oxidation and protein N-terminal acetylation were specified as variable modifications. MS2 spectra were searched against the *D. melanogaster* 1 protein per gene reference proteome from Uniprot (Proteome ID: UP000000803, release 2023.03), concatenated with a database of 382 common laboratory contaminants (release 2023.03, https://github.com/maxperutzlabs-ms/perutz-ms-contaminants).

Computational analysis was performed using Python and the in-house developed Python library MsReport (version 0.0.23). Only non-contaminant proteins identified with a minimum of two peptides and being quantified in at least two replicates of one experiment were considered for the analysis. LFQ protein intensities reported by FragPipe were log2-transformed and normalized across samples using the ModeNormalizer from MsReport. This method involves calculating log2 protein ratios for all pairs of samples and determining normalization factors based on the modes of all ratio distributions. For the data presented in the figures, missing values were imputed by deterministic lowest of detection (LOD) after filtering out contaminants and proteins with less than two razor and unique peptides, an approach we previously found to yield superior results for centrosomal proteins which are frequently absent in one or all control samples but also not highly abundant in the experimental samples (*36*). Also included in Data S2 is the more standard approach of imputation by drawing random values from a left-censored normal distribution modeled on the whole dataset (data mean shifted by −1.8 standard deviations, width of distribution of 0.3 standard deviations). Likely proximity interactors largely passed the significance threshold with both methods of imputation. These were defined as a log2 fold change of >1 and a p-value in an unpaired t-test of <0.05. GraphPad Prism was used to prepare bar graphs and volcano plots using LFQ values imported from Microsoft Excel.

### Comparisons to published human datasets and Gene Ontology (GO) analysis

To compare the proteins significantly enriched in our Cmb proximity interactome analyses in flies with those reported in published datasets of centrosomal, ciliary and centriolar satellite proteins as well as cytosolic mRNA-associated proteins in vertebrates (*3–5, 39*), we first generated a proteome-wide matrix of all conserved proteins based on reciprocal BLAST analysis as described above, then performed a pairwise comparison between datasets using Flybase IDs in *Drosophila* and Ensembl gene IDs in humans. Venn diagrams presented in the figures report both the number of conserved proteins common to both datasets and those unique to each, as well the larger number of unique proteins including those proteins not detectably conserved across species.

Gene Ontology analyses were performed using the PANTHER 18.0 (*95*) and GO Ontology database release 2023-10-09, comparing the protein IDs of the orthologs of proteins enriched in our Cmb proximity interactome analyses against the human proteome, identifying GO annotation terms statistically enriched based on Fisher’s Exact Test, applying Bonferroni correction for multiple testing. Annotation terms displayed in the figures are the top 8 terms for ‘cellular compartment’ and ‘biological process’ based on fold enrichment.

### Statistical analysis

Statistical analysis was performed in Excel and GraphPad Prism 10. Each dataset was tested for normal distribution using D’Agostino-Pearson (omnibus K2) test. Where data was displaying a normal distribution, an unpaired, two-tailed Student’s t-test with Welch’s correction was used for comparison of two groups. When data failed the normality test for at least one of the datasets examined, an unpaired, two-tailed Mann-Whitney U-test and a Kruskal-Wallis test with Dunn’s multiple comparisons test was used for two groups or more than two groups, respectively. If n was too small to test data for normal distribution a Student’s t-test was used. For all experiments the significance threshold was taken as P<0.05. Significance levels are defined as follows: ****P< 0.0001, ***P< 0.001, **P< 0.01, *P< 0.05, NS, not significant. Error bars display mean with standard deviation unless otherwise stated.

**Fig. S1.**
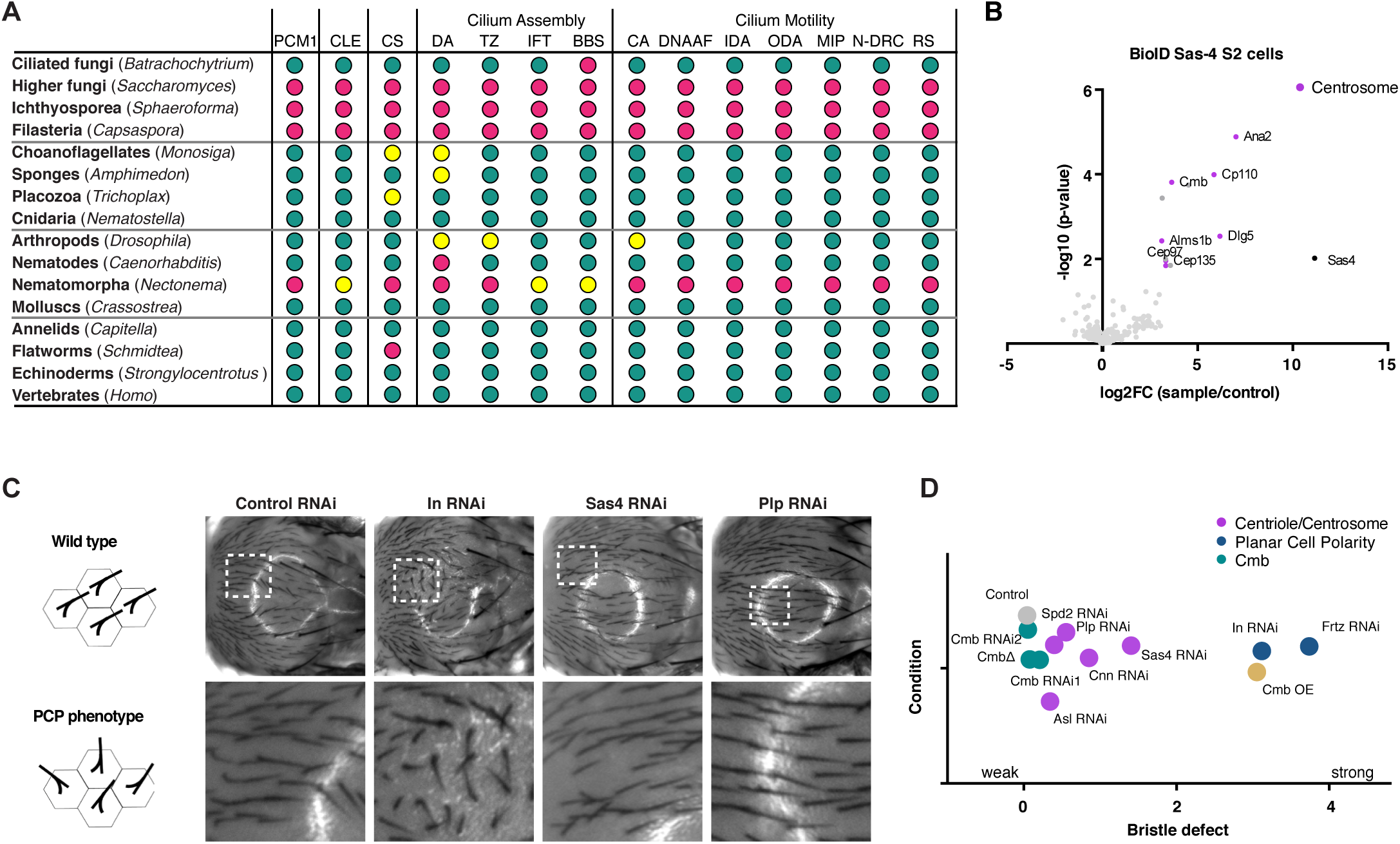
Identification of Combover as the *Drosophila* ortholog of PCM1. (**A**) Related to Fig. 1B. Conservation of PCM1 as well as core centriolar (STIL/Ana2, SASS6/Sas6, CENPJ/Sas4, CEP135/Bld10), centrosomal (CDK5RAP2/Cnn, CEP192/Spd2) and ciliary proteins (transition zone, IFT and BBS components, inner and outer dynein arm components, dynein assembly factors, nexins, N-DRC, radial spoke and central apparatus components, (*28*)) across opisthokonts, based on reciprocal BLAST analysis and hidden Markov model-based searches. Color code is green > 2/3 of genes in indicated category present, yellow > 1/3 of genes present, magenta < 1/3 present. See also Data S1. (**B**) Results of LC-MS/MS analysis for direct BioID performed on the centriolar structural component Sas4 in *Drosophila* S2 cells. Volcano plot of −log10 p-values against log2 fold change (sample/control). Significantly enriched proteins (log2 enrichment >1, p-value <0.05) indicated in dark grey, with centrosomal proteins highlighted in magenta. Cmb was detected as a high confidence interactor. See also Data S2A. (**C**) Related to Fig. 1D. Further characterization of PCP phenotypes in the fly notum. RNAi of PCP genes such as Inturned results in strong phenotypes, while centrosomal genes (Sas4 and Plp) show no or weak phenotypes. (**D**) Quantitation of bristle defects for selected genes. Phenotypes scored on a scale from 0 (no phenotype) to 4 (strong phenotype), with values shifted slightly to avoid overlap. N=10 flies per condition.

**Fig. S2.**
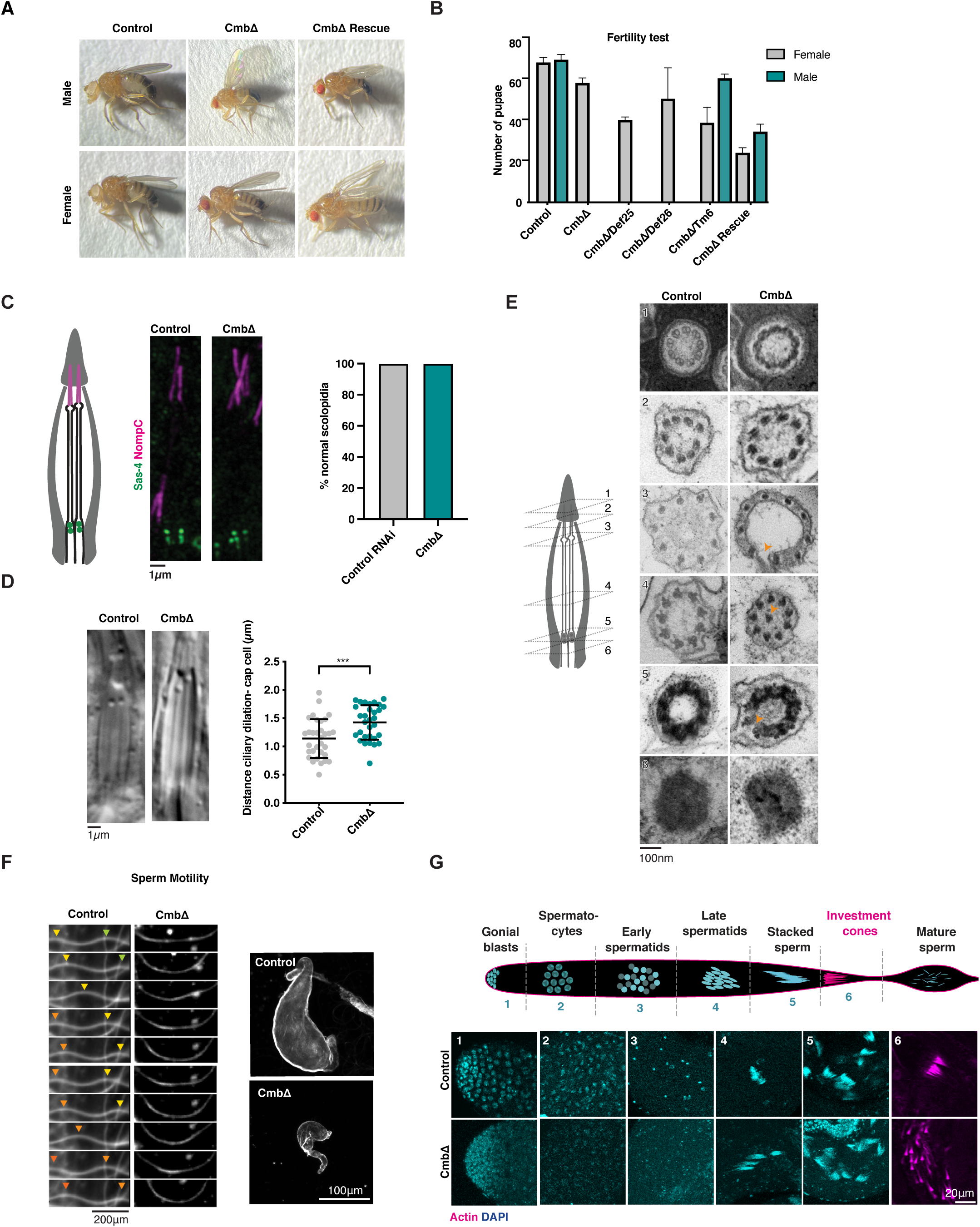
Further characterization of ciliary phenotypes in Cmb mutants. (**A**) Appearance of Cmb mutant flies, as well as Cmb mutants rescued by expression of a GFP-Cmb transgene. Cmb mutants display abnormal wing posture, a phenotype associated with defective mechanosensation. (**B**) Fertility test performed on Cmb mutant males and females, Cmb mutants rescued by expression of a GFP-Cmb transgene or maintained over a balancer (Tm6) and Cmb mutants placed over a deficiency that covers the Cmb locus (Def 25, 26). Cmb mutant males but not females exhibit fully penetrant sterility, a defect rescued by expression of the GFP transgene. Placing the mutant over a deficiency does not impact fertility, excluding potential non-allelic effects. N=3 single males, each crossed to 4 virgin females. (**C**) Schematic and immunofluorescence micrographs of scolopidia in chordotonal organ of the fly. Sas4 and NompC used to visualize basal body (green) and ciliary tip (magenta), respectively. Each scolopidium contains two cilia with their ends embedded in the cap cell. No apparent ciliary structural defects are observed in Cmb mutants. N=63 control scolopidia, 63 Cmb mutant. Statistical test is t-test with Welch’s correction. (**D**) DIC images of scolopidia. Cmb mutants display a larger distance between ciliary dilation and cap cell, indicative of ciliary positioning defects. N=32 control, N=31 Cmb mutants. Student’s t-test with Welch’s correction used; ***P< 0.001. (**E**) Cross sectional views of control and Cmb mutant scolopidia by transmission electron microscopy (TEM). Position indicated by numbers in schematic on the left. Cmb mutants show minor structural defects, including broken axonemes and misplaced doublet microtubules. (**F**) Left: Analysis of flagellar movement of control and Cmb mutant sperm by high-speed video capture in dark-field microscopy. Sinusoidal motion can be seen in wild type. Cmb mutant sperm show severely compromised flagellar movement. Right: In contrast to controls, seminal vesicles of Cmb mutant flies are almost devoid of sperm, indicating defective movement of sperm to seminal vesicle. (**G**) Schematic and immunofluorescence images of *Drosophila* spermatogenesis. In Cmb mutants the early stages of spermatogenesis appear superficially normal; however, in later stages investment cones involved in individualizing sperm fail to form properly.

**Fig. S3.**
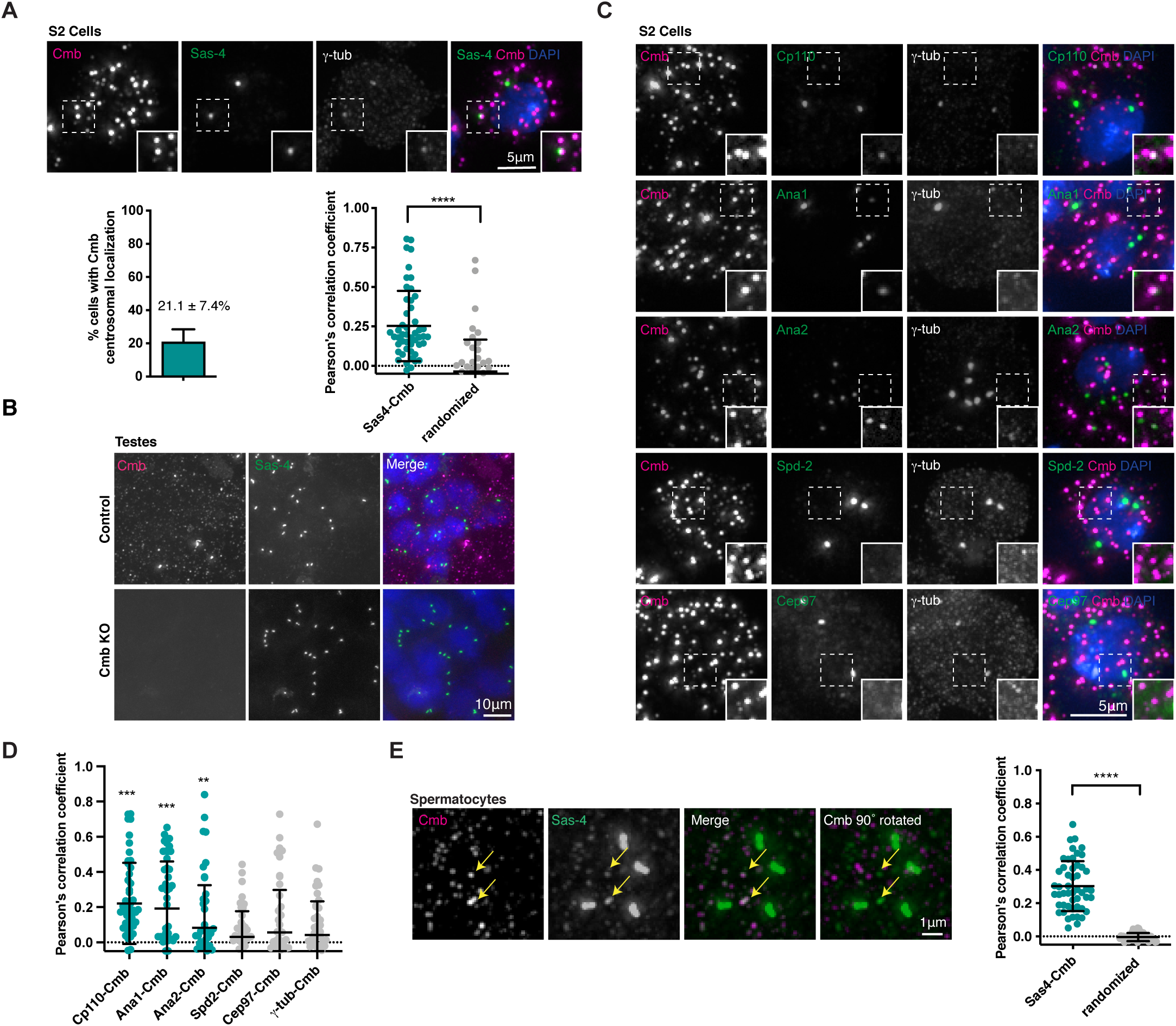
Further analysis of Cmb localization. (**A**) Immunofluorescence micrograph showing Cmb localizing to centrosomes in a subset of cells in S2 cells. Quantitation of Cmb localization reveals centrosome localization in ∼20% of cells. Pearson correlation coefficient analysis of centrosomal signal shows significant overlap between Sas-4 and Cmb compared to randomized controls (single channel rotated by 90° with centrosome positioned in the upper right quadrant of the square analyzed). Student’s t-test was used; ****P< 0.0001. N=50 cells. (**B**) Specificity of polyclonal antibody raised against Cmb confirmed by absence of immunofluorescence signal in Cmb mutant testes. (**C**) Related to Fig. 3I) Immunofluorescence micrographs showing some (Cp110, Ana1, Ana2) but not all (Spd-2, γ-tubulin) centrosomal proteins colocalizing with Cmb on cytoplasmic foci. Pearson’s correlation coefficient analysis on cytoplasmic signal assessed as in Fig. 3G. N=50 cells per condition. Mann-Whitney test used to test statistical significance; ***P< 0.001, **P< 0.01. (**D**) Immunofluorescence micrographs showing Sas4 colocalizing with Combover in the cytoplasm of primary spermatocytes in the testes. Colocalization quantified by Pearson’s correlation coefficient analysis. N=12 animals. t-test used to assess statistical significance of colocalization compared to randomized controls (single channel rotated 90°); ****P< 0.0001.

**Fig. S4.**
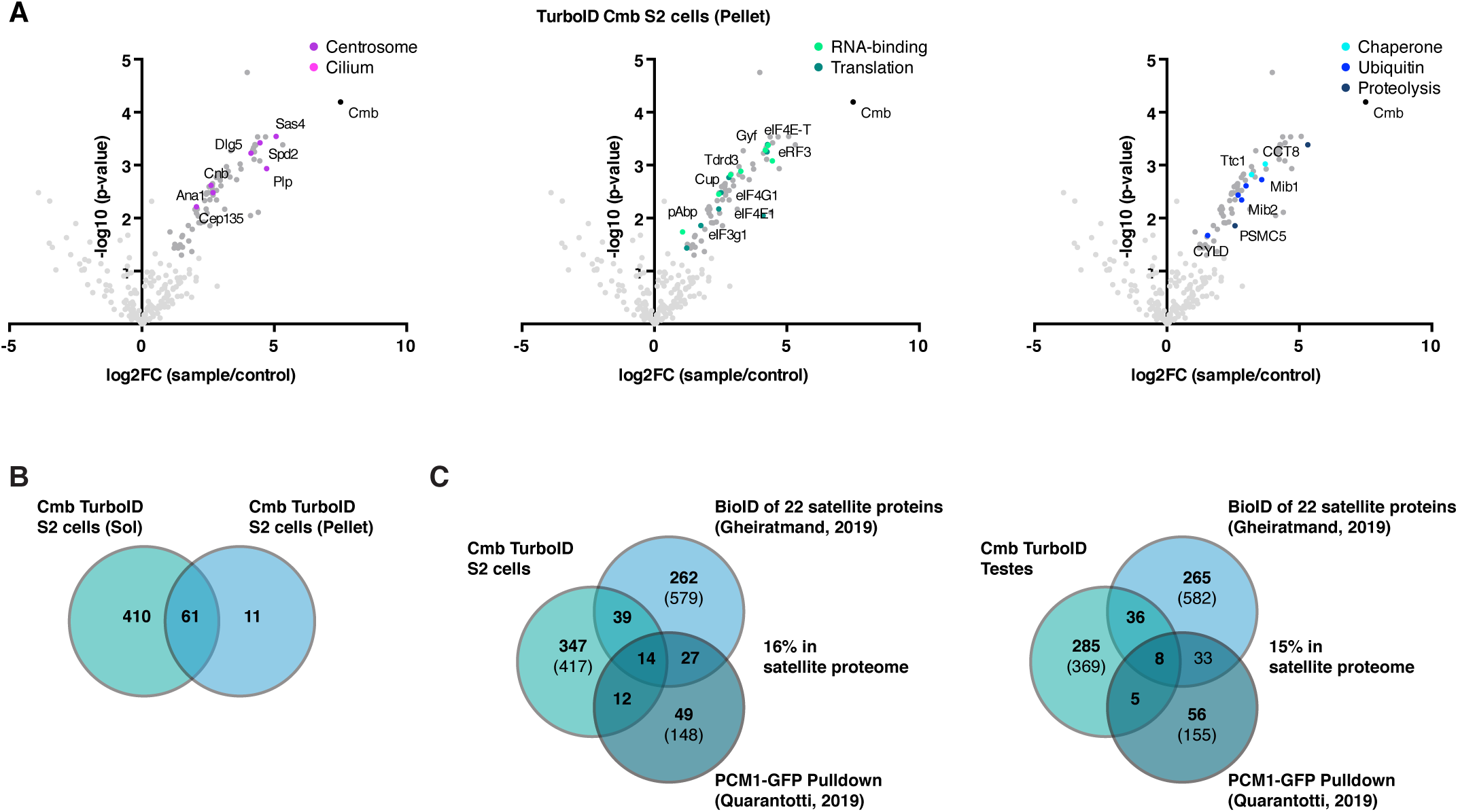
Further analysis of Cmb proximity interactome. (**A**) Related to Fig. 4A. Results of direct TurboID performed on Cmb S2 cells (detergent-insoluble cytoskeletal fraction). LC-MS/MS analysis reveals centrosomal proteins (magenta), RNA-binding proteins (light green) and proteins involved in translation (dark green), chaperone-mediated protein folding (light blue), ubiquitination (blue) and proteolysis (dark blue). Volcano plots of −log10 p-values against log2 fold change (sample/control). Significantly enriched proteins (Log2 enrichment >1, p-value <0.05) indicated in dark grey, with proteins of the above functional categories highlighted in color. See also Data S2C. (**B**) Venn diagram revealing significant overlap between Cmb proximity interactome obtained from detergent-soluble (cytoplasmic) and detergent-insoluble (cytoskeletal) fractions of S2 cell extracts. See also Data S2E. (**C**) Comparison of Cmb S2 cell and testes TurboID interactomes with previous published datasets for centriolar satellites: the BioID of 22 satellite proteins mapped by (*4*) and the PCM1-GFP pulldown performed by (*5*). Comparison for those proteins conserved between human and flies. Numbers in parentheses are total number in each dataset. See also Data S2E.

**Fig. S5.**
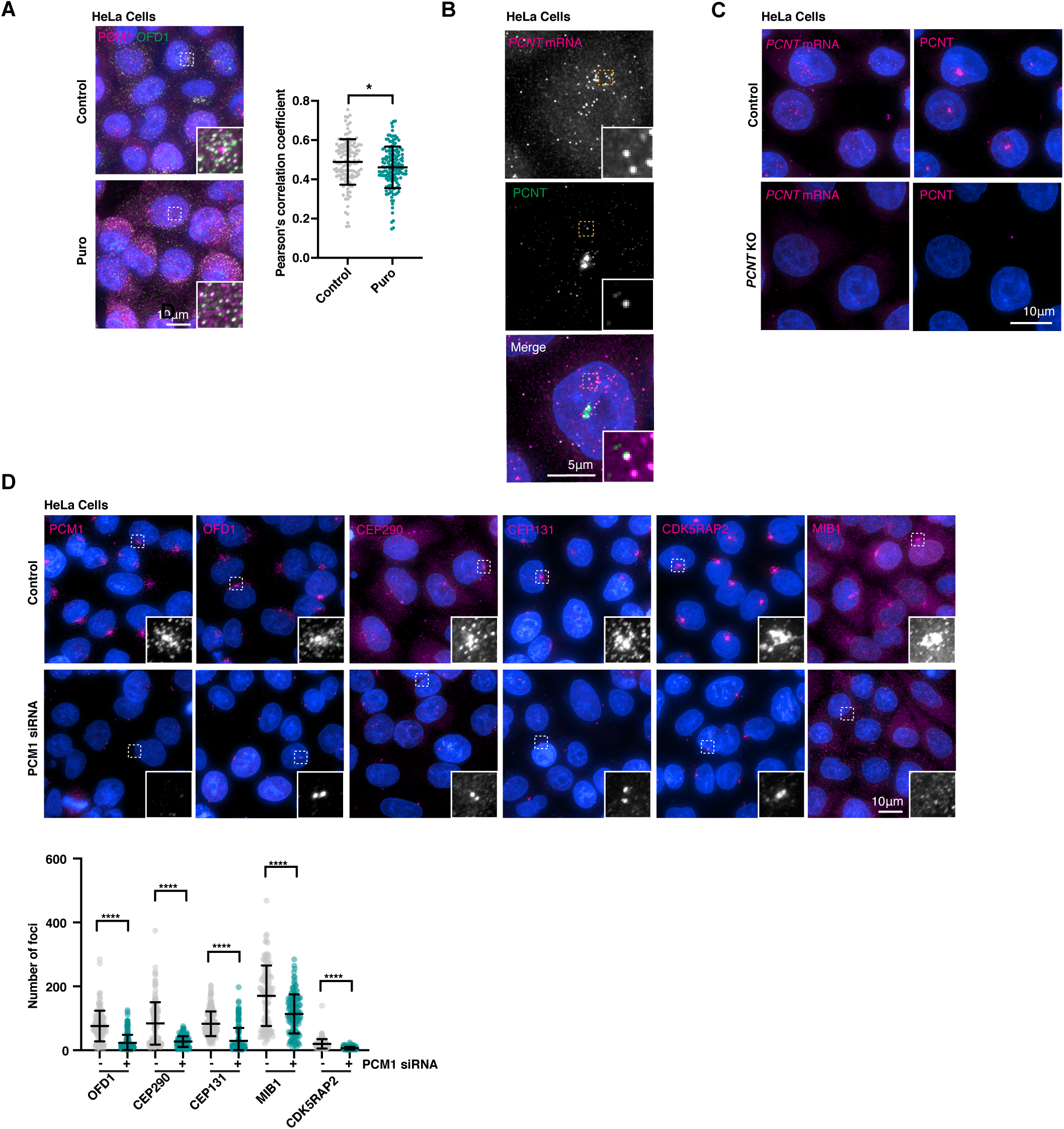
Further evidence for centriolar satellites as sites of translation in vertebrate cells. (**A**) Immunofluorescence micrographs and quantitation of PCM1 and OFD1 distribution by Pearson’s correlation coefficient analysis in control and puromycin-treated cells. PCM1 and OFD1 co-localize independent of translation. N=125 cells control, 147 puromycin treatment. Mean +/-SD indicated (Students t-test; *P<0.05). (**B**) SmFISH combined with immunofluorescence microscopy shows pcnt mRNA colocalizing with nascent Pcnt protein in the cytoplasm. (**C**) Specificity of pcnt mRNA FISH and protein immunofluorescence signal confirmed by absence of signal in *PCNT* KO cells. (**D**) Immunofluorescence micrographs and quantitation of centriolar satellite signal in control and PCM1 siRNA treated cells. Depletion of PCM1 largely eliminates cytoplasmic foci of OFD1, CEP290, CEP131, CDK5RAP2, while centrosomal signal remains. MIB1 signal is largely unaffected, although foci are now dispersed throughout the cytoplasm. Mean +/-SD indicated. N>100 cells each condition. Statistical test to compare control and PCM1 depletions is t-test with Welch’s correction (MIB1), non-parametric Mann Whitney test (others); ****P<0.0001.

**Table S1.** Fly strains used in this study.

| Strain description | Source | Identifier |
| --- | --- | --- |
| Asl RNAi/ P{TRiP.HMS01453}attP2 | BDSC | BDSC ID: 35039, RRID: BDSC_35039 |
| Asl-GFP & H2A-RFP; CmbΔ/ P{His2Av-mRFP1}; P{UAS.asl-asl-GFP.B}; CmbΔ/Tmb6 SbTb | This study | N/A |
| Asl-GFP & H2A-RFP/ P{His2Av-mRFP1}; P{UAS.asl-asl-GFP.B} | This study | N/A |
| Asl-GFP/ P{UAS.asl-asl-GFP.B} | (64) | Flybase ID: FBtp0040947 |
| Bam-GAL4 & Nanos-GAL4/ P{GAL4-nos.NGT}40; P{bam-GAL4:VP16} | (28) | N/A |
| Cmb OE/ P{UAS-cmb.RA} | (23) | Flybase ID: FBtp0095481 |
| Cmb RNAi 1/ P{KK103563}VIE-260B | VDRC | VDRC ID: 109767, RRID: Flybase_FBst0481420 |
| Cmb RNAi 2/ P{GD7077}v18666 | VDRC | VDRC ID: 18666, RRID: Flybase_FBst0453208 |
| Cmb-GFP rescue/ PBac{fTRG01063.sfGFP-TVPTBF}VK00002; CmbΔ | This study | N/A |
| Cmb-GFP/ PBac{fTRG01063.sfGFP-TVPTBF}VK00002 | VDRC | VDRC ID: 318739, RRID: Flybase_FBst0491914 |
| CmbΔ/ CmbΔ/Tm6 SbTb | (23) | Flybase ID: FBal0298847 |
| Cnn RNAi/ P{GD1726}v44526 | VDRC | VDRC ID: 44526, RRID: Flybase_FBst0465625 |
| Control RNAi/ P{TRiP.HMS00045}attP2 | BDSC | BDSC ID: 33644, RRID: BDSC_33644 |
| Deficiency 1/ w[1118]; Df(3L)BSC614/TM6C, cu[1] Sb[1] | BDSC | BDSC ID: 25689, RRID: BDSC_25689 |
| Deficiency 2/ w[1118]; Df(3L)BSC737/TM6C, Sb[1] cu[1] | BDSC | BDSC ID: 26835, RRID: BDSC_26835 |
| Elav-GAL4/ P{GawB}elav[C155] | BDSC | BDSC ID: 458, RRID: BDSC_458 |
| Frtz RNAi/ P{GD9437}v40088 | VDRC | VDRC ID: 40088, RRID: Flybase_FBst0463379 |
| GFP nanobody-TurboID/ P{egg-TurboID.vhhGFP.3xHA} | This study | N/A |
| H2A-RFP/ P{His2Av-mRFP1} P{UAS.asl-asl-GFP.B} | (63) | Flybase ID: FBtp0056035 |
| In RNAi/ P{GD10892}v27252 | VDRC | VDRC ID: 27252, RRID: Flybase_FBst0456836 |
| Plp RNAi/ P{GD9625}v20667 | VDRC | VDRC ID: 20667 |
| Pnr-GAL4/ P{GawB}pnr[MD237] | (66) | Flybase ID: FBti0004011 |
| RFP-Cnn; CmbΔ/ P{Ubi-p63E-cnn.RFP};<br>CmbΔ/Tmb6 SbTb | This study | N/A |
| RFP-Cnn/ P{Ubi-p63E-cnn.RFP} | (62) | Flybase ID: FBtp0056991 |
| Sas4 RNAi/ P{GD6852}v17975 | VDRC | VDRC ID: 17975, RRID:<br>Flybase_FBst0452943 |
| Spd2 RNAi/ P{KK110116}VIE-260B | VDRC | VDRC ID: 101882, RRID:<br>Flybase_FBst0473755 |
| Wild-type/ w[1118] | BDSC | BDSC ID: 6326, RRID:<br>BDSC_6326 |

**Table S2.**
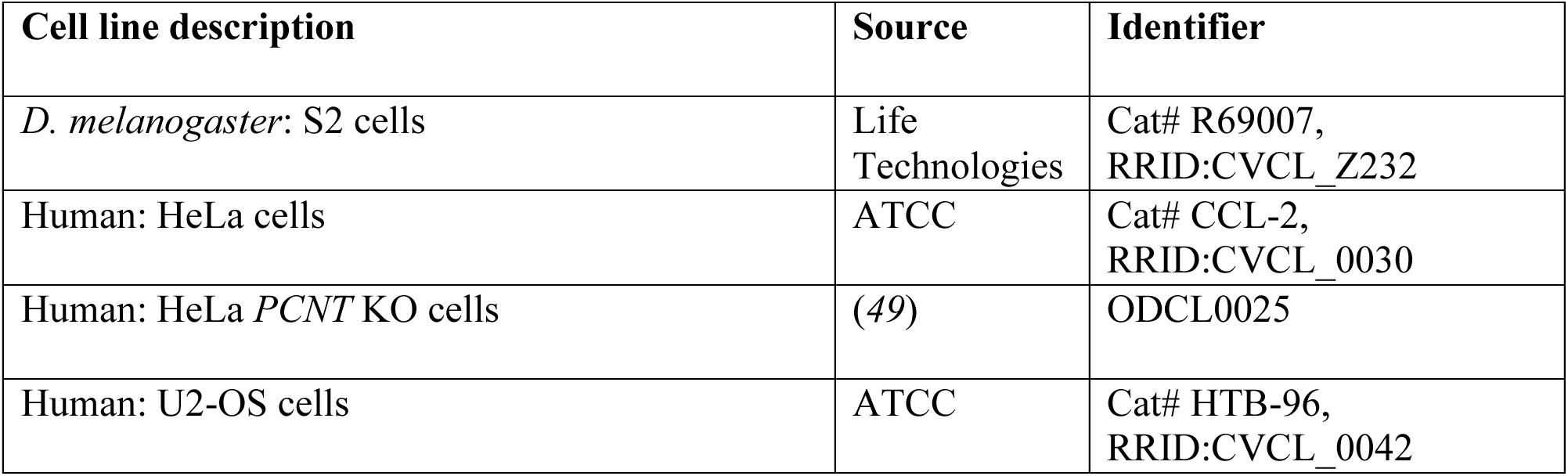
Cell lines used in this study.

| Cell line description | Source | Identifier |
| --- | --- | --- |
| *D. melanogaster*: S2 cells | Life Technologies | Cat# R69007,<br>RRID:CVCL_Z232 |
| Human: HeLa cells | ATCC | Cat# CCL-2,<br>RRID:CVCL_0030 |
| Human: HeLa *PCNT* KO cells | (49) | ODCL0025 |
| Human: U2-OS cells | ATCC | Cat# HTB-96,<br>RRID:CVCL_0042 |

**Table S3.** Plasmid constructs used in this study.

| **Plasmid description** | **Source** | **Identifier** |
| --- | --- | --- |
| Blasticidin selection vector: pCoBlast | Thermo Fisher | Cat# K515001 |
| GST expression vector: pGEX-6P-1 Cmb (aa1-150) | This study | N/A |
| BioID control vector: pMt-myc-BirA* | This study | N/A |
| BioID Sas4 vector: pMt-Sas4-myc-BirA* | This study | N/A |
| S2 cell expression vector: pMt-V5-6xHis B | Invitrogen | Cat# V412020 |
| Cmb GFP vector: pMt-V5-6xHis-Cmb-GFP | This study | N/A |
| TurboID Cmb vector: pMt-V5-TurboID-Cmb | This study | N/A |
| TurboID control vector: pMt-V5-TurboID-GW | Norbert Perrimon | N/A |
| Indirect TurboID fly expression vector: pUASz-TurboID-vhhGFP-3xHA | This study | N/A |
| Indirect TurboID expression vector (+NLS): pUASz-TurboID-vhhGFP-3xHA-NLS | (67) | N/A |
| Gateway vector: pDONR Zeo | Thermo Fisher | Cat# 12535035 |
| BioID source vector: pcDNA3.1 mycBirA* | Kyle Roux | N/A |

**Table S4.** Antibodies used in this study.

| **Antibody description** | **Source** | **Identifier** |
| --- | --- | --- |
| Donkey anti-Mouse IgG (H + L) Cy2-conjugated antibody | Jackson ImmunoResearch | Cat# 715-225-150, RRID: AB_2340826 |
| Donkey anti-Mouse IgG (H + L) Cy3-conjugated antibody | Jackson ImmunoResearch | Cat# 715-165-150, RRID: AB_2340813 |
| Donkey anti-Mouse IgG (H + L) Cy5-conjugated antibody | Jackson ImmunoResearch | Cat# 715-175-150, RRID: AB_2340819 |
| Donkey anti-Rabbit IgG (H + L) Cy2-conjugated antibody | Jackson ImmunoResearch | Cat# 711-225-152, RRID: AB_2340612 |
| Donkey anti-Rabbit IgG (H + L) Cy3-conjugated antibody | Jackson ImmunoResearch | Cat# 711-165-152, RRID:AB_2307443 |
| Donkey anti-Rabbit IgG (H + L) Cy5-conjugated antibody | Jackson ImmunoResearch | Cat# 711-175-152, RRID:AB_2340607 |
| Mouse monoclonal anti-dmNompC antibody | (74) | N/A |
| Mouse monoclonal anti-gamma Tubulin antibody | Sigma-Aldrich | Cat# T6557, RRID: AB_477584 |
| Mouse monoclonal anti-hsPCM1 antibody | Santa Cruz Biotechnology | Cat# sc-398365, RRID: AB_2827155 |
| Mouse monoclonal anti-Puromycin | Merck Millipore | Cat# MABE343, RRID: AB_2566826 |
| Rabbit polyclonal anti-dmAna1 antibody | (62) | N/A |
| Rabbit polyclonal anti-dmAna2 antibody | (77) | N/A |
| Rabbit polyclonal anti-dmCep97-N-ter antibody | (27) | N/A |
| Rabbit polyclonal anti-dmCmb antibody | This study | N/A |
| Rabbit polyclonal anti-dmCnn antibody | (75) | N/A |
| Rabbit polyclonal anti-dmCP110 antibody | (76) | N/A |
| Rabbit polyclonal anti-dmSas4 antibody | (30) | N/A |
| Rabbit polyclonal anti-dmSpd2 antibody | (77) | N/A |
| Rabbit polyclonal anti-hsCDK5RAP2 antibody | Merck Millipore | Cat# 06-1398, RRID: AB_11203651 |
| Rabbit polyclonal anti-hsCEP131/AZI1 antibody | Abcam | Cat# ab84864, RRID: AB_1859791 |
| Rabbit polyclonal anti-hsCEP290 antibody | Novus Biologicals | Cat# NB100-86991, RRID: AB_1201171 |
| Rabbit polyclonal anti-hsMIB1 antibody | Sigma-Aldrich | Cat# M5948; RRID: AB_1841007 |
| Rabbit polyclonal anti-hsOFD1 antibody | Sigma-Aldrich | Cat# HPA031103, RRID: AB_10602188 |
| Rabbit polyclonal anti-hsPCM1 antibody | (6) | N/A |
| Rabbit polyclonal anti-hsPCNT antibody | Abcam | Cat# ab4448, RRID: AB_304461 |

**Table S5.** Chemicals, enzymes and other reagents used in this study.

| Reagent description | Source | Identifier |
| --- | --- | --- |
| Agar 100 Resin | Agar Scientific | Cat# AGR1031; CAS: 25038–04-4 |
| Cycloheximide | Sigma-Aldrich | Cat# C1988; CAS: 66-81-9 |
| 25% Glutaraldehyde, EM grade | Agar Scientific | Cat# AGR1020; CAS: 111-30-8 |
| Hoechst 33258 | Thermo Fisher | Cat# H1398; CAS: 23491–45-4 |
| Lipofectamine RNAiMax | Invitrogen | Cat# 13778100 |
| Lysyl Endopeptidase (LysC) | FUJIFILM Wako chemicals | Cat# 129-02541 |
| Osmium Tetroxide 4% Solution | Electron Microscopy Sciences | Cat# 19170; CAS: 20816–12-0 |
| Paraformaldehyde | Roth | Cat# O335.1; CAS: 30525-89-4 |
| 32% Paraformaldehyde, EM grade | Electron Microscopy Sciences | Cat # 15714 |
| Phalloidin Alexa Fluor 568 | Thermo Fisher | Cat# A12380 |
| Pierce Streptavidin Magnetic Beads | Thermo Scientific Pierce | Cat# 88817 |
| Puromycin dihydrochloride | Sigma-Aldrich | Cat# P8833 |
| Reynolds lead citrate | Delta Microscopies | Cat# 11300; CAS: 512–26-5 |
| TransIT Insect Transfection reagent | Mirus Bio | Cat# MIR 6104 |
| Trypsin Gold Mass Spec Grade | Promega | Cat# V5280 |
| Uranyl acetate | Science Services | Cat# E22400; CAS: 541–09-3 |
| Vanadyl ribonucleoside complexes solution | Sigma-Aldrich | Cat# 94742 |
| Vectashield Plus | Vector Laboratories | Cat# H-1900 |

**Table S6.**
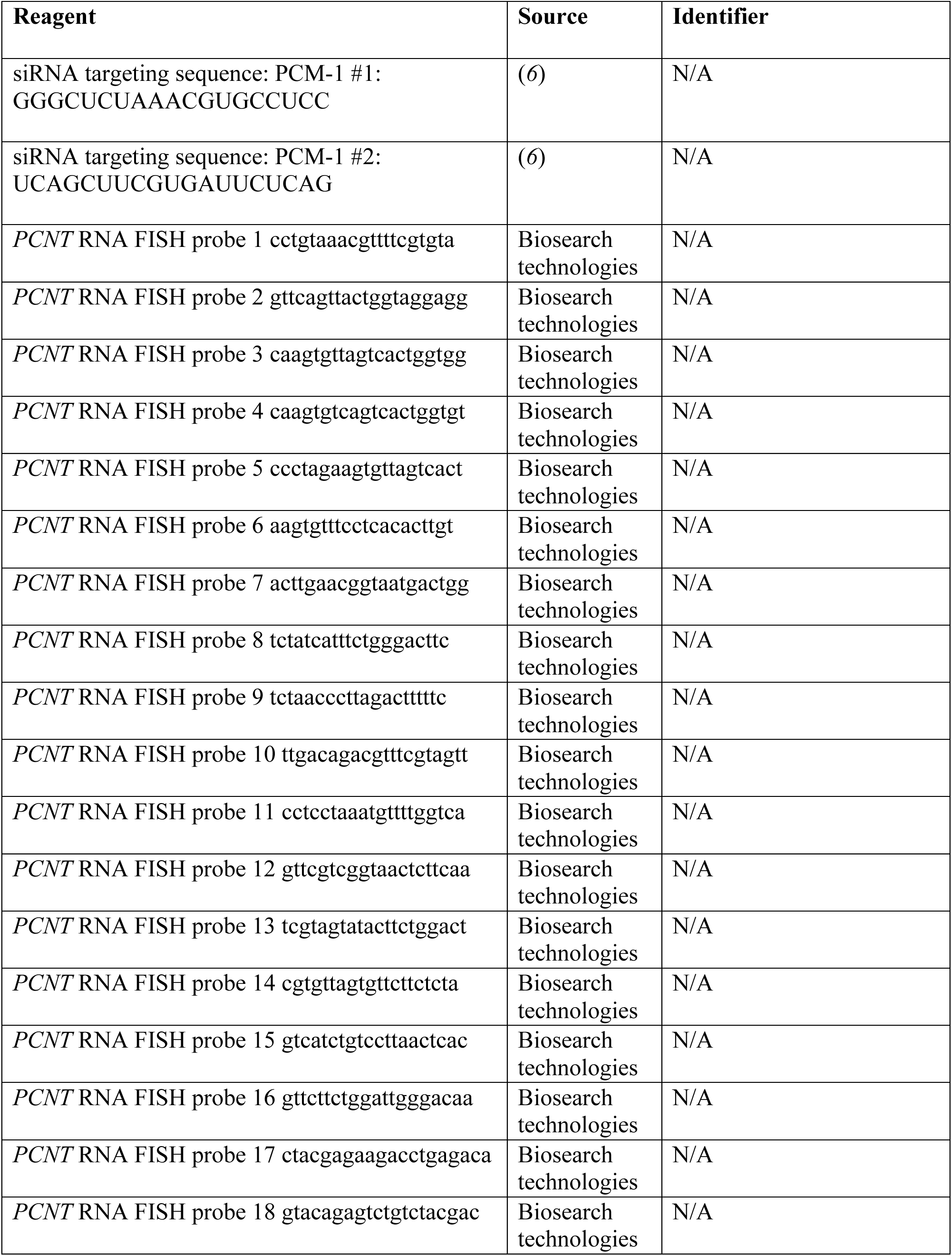
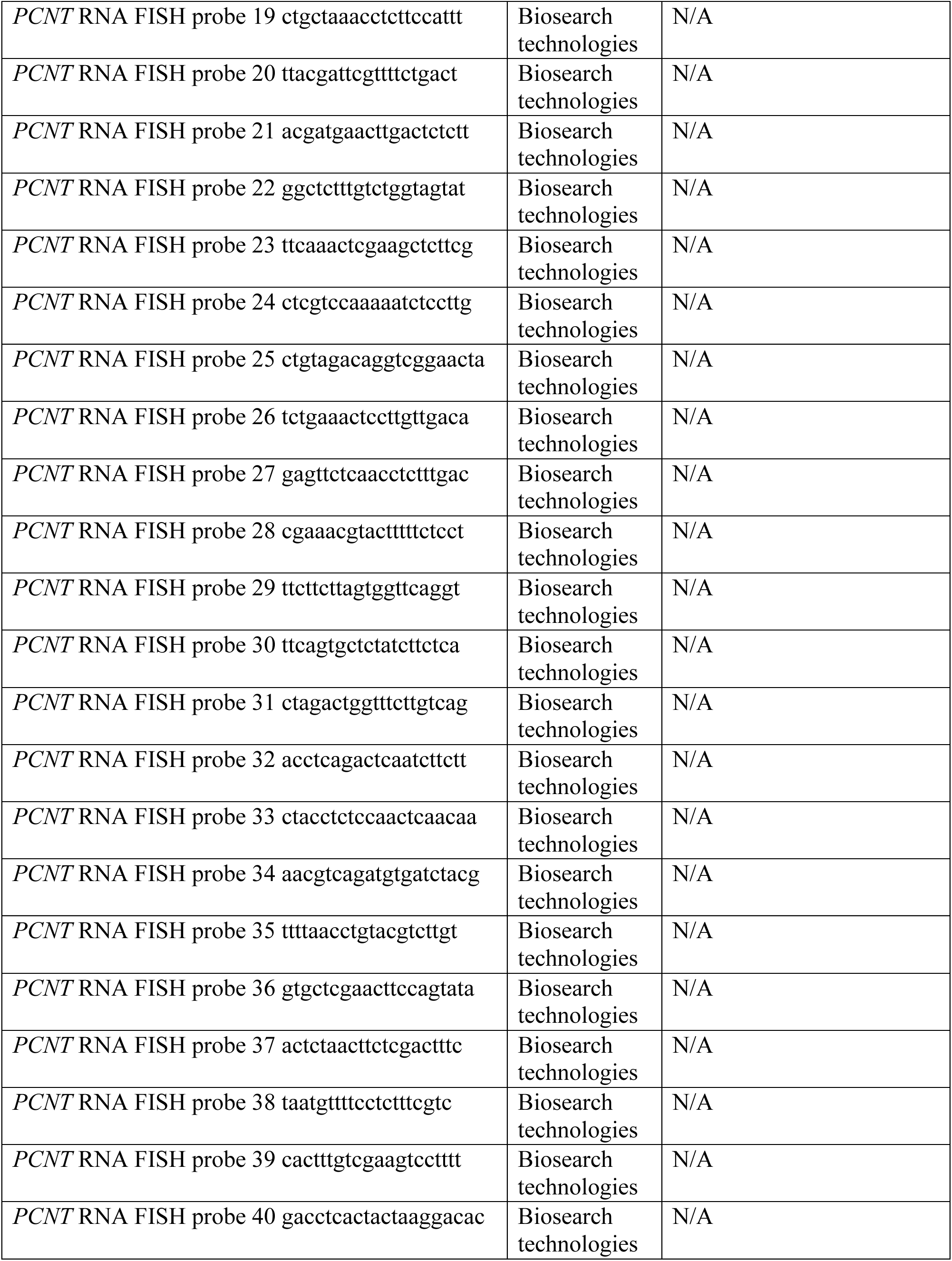
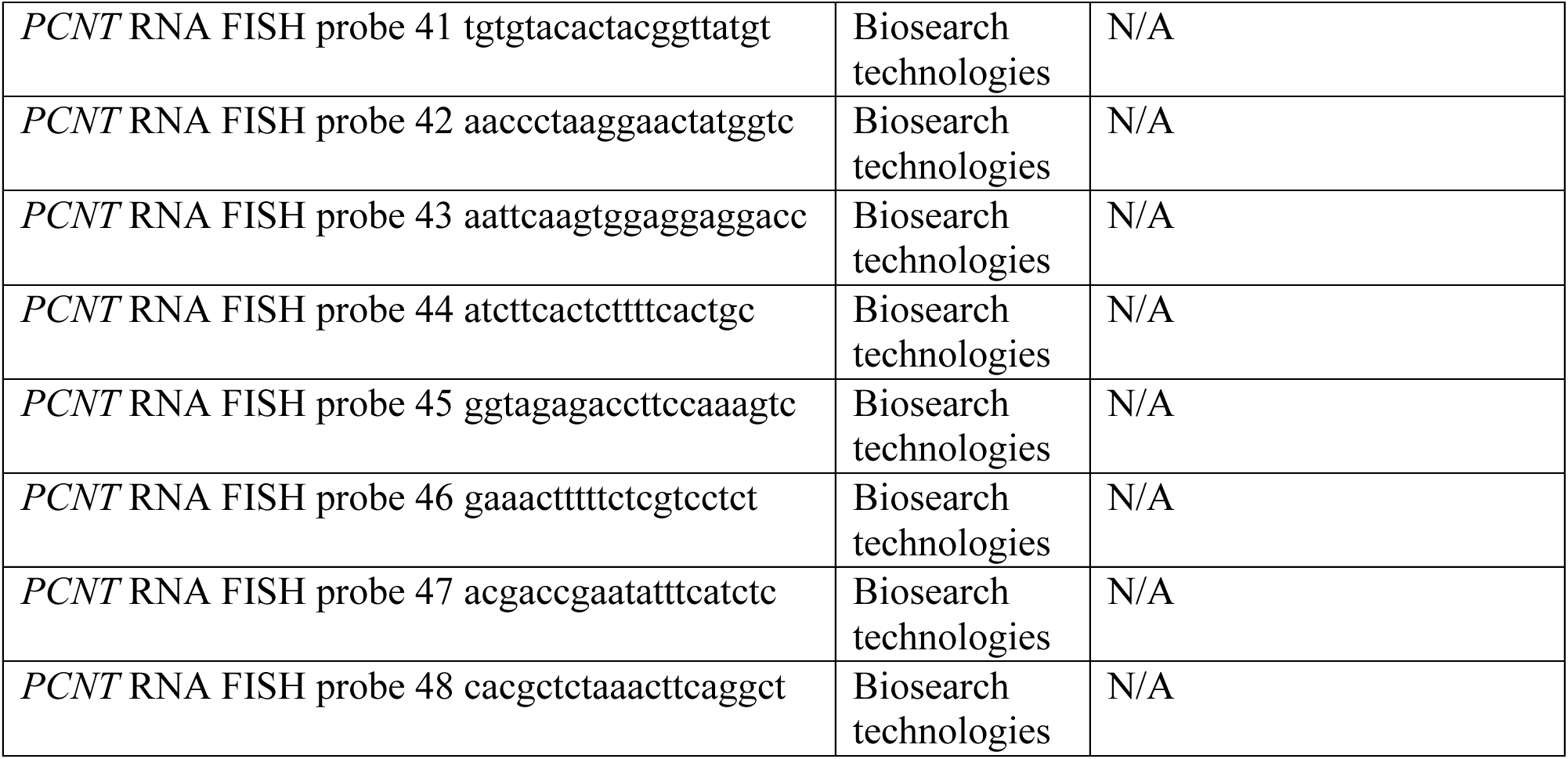
Oligonucleotides used in this study.

**Table S7.** Software and algorithms used in this study.

| Software description | Source | Identifier |
| --- | --- | --- |
| Fiji v 2.0.0 | NIH | https://imagej.net/software/fiji/ |
| FragPipe v 20.0 | University of Michigan Medical School | https://github.com/Nesvilab/FragPipe |
| GraphPad Prism 10 | GraphPad | https://www.graphpad.com/scientific-software/prism |
| HMMER v 3.4 | (71) | http://hmmer.org/ |
| Ilastik v 1.4.0 | (85) | https://www.ilastik.org |
| IonQuant v 1.9.8 | (93) | https://github.com/Nesvilab/IonQuant |
| Jalview v 2.11.1.4 | (96) | https://www.jalview.org/ |
| MAFFT v 7 | (72) | https://mafft.cbrc.jp/alignment/server/ |
| Matlab v R2023a | MathWorks | https://www.mathworks.com |
| MSFragger v 3.8 | (92) | https://github.com/Nesvilab/MSFragger |
| MsReport v 0.0.23 | Max Perutz Labs Mass Spectrometry Facility | N/A |
| NCBI BLAST+ v 2.13.0 | (97) | ftp://ftp.ncbi.nlm.nih.gov/blast/executables/blast+/LATEST/ |
| Philosopher v 5.0.0 | (94) | https://github.com/Nesvilab/philosopher |

**Data S1. Orthologs of PCM1 and centrosomal/ciliary proteins across Opisthokonts.** (**A**). Orthologs of PCM1, CDK5RAP2 and CEP192 in representative species. Related to Fig. 1B. (**B**) Orthologs of core centriolar (STIL/Ana2, SASS6/Sas6, CENPJ/Sas4, CEP135/Bld10) and ciliary proteins (transition zone, IFT and BBS components, inner and outer dynein arm components, dynein assembly factors, nexins, N-DRC, radial spoke and central apparatus components, (*28*)) in Nematomorpha, based on reciprocal BLAST analysis. Related to Figs. 1B and S1A. (**C**) GenBank accession numbers for PCM1 orthologs used to generate multiple sequence alignment in Fig. 1C.

**Data S2. Mass spectrometry data. (A**-**D)** Complete list of proteins identified by mass spectrometry in each BioID/TurboID run, including corresponding controls. (A) Sas4 BioID S2 cells. (B) Cmb TurboID S2 cells soluble/cytoplasmic fraction. (C) Cmb TurboID S2 cells insoluble/cytoskeletal fraction. (D) Cmb TurboID testes. Only the most relevant data rows and columns are displayed by default. The full processed MS data including excluded peptide groups, spectral counts and differential abundance analysis using missing value imputation by random drawing from a left-censored normal distribution (ND) can be found by expanding the collapsed rows and columns. (**E**) Comparison of Cmb TurboID proximity interactors with published datasets of centrosomal, ciliary and centriolar satellite proteins as well as cytosolic mRNA-associated proteins in vertebrates (*3–5, 39*).

